# Comparative pan-genomics reveals extensive variation in secondary metabolism and the non-coding repertoire of clinically-relevant *Fusarium solani* species complex members

**DOI:** 10.1101/2024.09.23.614396

**Authors:** Phillip J.T. Brassington, M. Laura Fabre, Anna Zimmermann, Lea M. Krämer, Marion Perrier, Annika Schöninger, Ronny Martin, Oliver Kurzai, Amelia E. Barber

**Author notes:** Address correspondence to: Amelia E. Barber.

## Abstract

The *Fusarium solani* species complex (FSSC) is a group of dual-kingdom fungal pathogens capable of causing devastating disease on a wide range of host plants and life-threatening infections in humans that are difficult to treat. In this study, we generate highly contiguous genomes for three clinical isolates of *Fusarium keratoplasticum* and three clinical isolates of *Fusarium petroliphilum* and compare them with other genomes of the FSSC from plant and animal sources. We find that human pathogenicity is polyphyletic within the FSSC, including within *F. keratoplasticum*. Pan-genome analysis revealed extensive gene presence-absence in the complex, with only 41% of genes (11,079/27,068) found in all samples and the presence of accessory chromosomes encoding isolate- and species-specific genes. Definition of long non-coding RNAs (lncRNAs) in *F. keratoplasticum* and *F. petroliphilum* demonstrated that they show a similarly high degree of presence-absence variation but are integrated into the transcriptional network to a similar degree as protein-coding genes, suggesting broad and extensive cellular roles. Elucidation of secondary metabolic potential identified many actively transcribed biosynthetic gene clusters with unknown products and identified several lncRNAs as putative regulators of specific metabolite clusters. Finally, examination of the FSSC for *Starship* gigantic mobile genetic elements revealed that *F. keratoplasticum* had the largest number with a median of four elements per genome and an overall pattern of Starships in the FSSC that were incongruent with the phylogeny, suggesting multiple horizontal transfer events. This study provides valuable insights into the evolutionary dynamics and genomic architecture of the FSSC, with implications for understanding multi-kingdom virulence, something of increasing relevance as climate change potentially increases the number of fungal species that can grow at human temperatures.

## Introduction

The genus *Fusarium* encompasses a highly diverse range of filamentous fungal species, including environmental saprobes, plant pathogens, and organisms capable of causing disease in animals and humans. Well established as plant pathogens, *Fusarium* spp. are causal agents for devastating worldwide food supply disruptions, causing significant losses in global yields of wheat, maize, rice, potato, and soybean, with outbreaks of *Fusarium* head blight causing over 50% wheat losses in China, Europe, and the Americas (1–5). Yet beyond this, *Fusarium* spp. are largely unique among fungi in that they are multi-kingdom pathogens capable of also infecting higher eukaryotes (6–8). In this capacity, *Fusarium* spp. play a role in the decline of threatened animal species (9–11), and are the causative agents with rising numbers of highly lethal human fusariosis infections (12). In 2022, the World Health Organization identified the genus as one of the highest priority human fungal pathogens, and with the most urgent need for additional study based on current knowledge gaps (13). These include lack of diagnostic options, limited epidemiological data, and intrinsic antifungal resistance, which cumulatively contribute to a high mortality rate of 40-70% (13–17).

Previous studies have highlighted the exceptional range of genomic diversity and plasticity among *Fusarium* spp. Even within a single species, the genome structure is highly flexible and strains often possess accessory chromosomes that are variably conserved and, importantly, confer the ability to infect specific plant species (18). These accessory chromosomes can even be horizontally transferred, resulting in a corresponding transfer of host plant specificity (19, 20). Despite the large number of species in the genus, the majority (reportedly up to 80-90%) of human infections are due to only two species groups: the *Fusarium oxysporum* species complex and the *Fusarium solani* species complex (8, 21–24).

The *Fusarium solani* species complex (FSSC) comprises globally ubiquitous problematic agricultural pathogens and is the most frequent cause of human fusariosis (8, 21, 24–26). Among the human pathologies it causes, it is particularly problematic in causing corneal infection and disseminated disease. More recently, it has been linked to two outbreaks of meningitis in immunocompetent patients in Mexico due to contaminated anesthetic solutions (27, 28). In the plant kingdom, the FSSC exhibits a broad host range and is capable of causing root rot in multiple agriculturally important crops, including peas, cucurbits, and sweet potato (29–32). Even within individual species of the complex, there is a broad range of habitats and hosts. For example, *F. keratoplasticum* has been found in diverse habitats ranging from plumbing systems, strawberry roots, Chinese softshell turtle eggs, to human nails, eyes, and endocardium (8, 11, 23, 33–35). This is also likely a non-exhaustive list, as many studies classify isolates as *forma specialis* (*f. sp.*). based on the host plant, rather than identifying the exact FSSC species member. The diversity of habitats and pathologies caused by the FSSC in both plants and animals is accompanied by a correspondingly high level of genomic diversity, including the presence of core and accessory chromosomes (11). However, the genomes of FSSC members remain understudied compared to other *Fusarium* groups like the *Fusarium oxysporum* or *graminearum* species complexes. Moreover, despite its relevance for human health, no FSSC isolates from human infections have been sequenced to date.

To address this, we defined the pan-genomic diversity of six FSSC member species, focusing on *F. keratoplasticum* and *F. petroliphilum*, two of the most prominent FSSC species in human keratitis (8, 36). We generated near-chromosomal genomes for the first clinical isolates of *F. keratoplasticum* and provide the first genomes to date for *F. petroliphilum*. We integrate these six clinical genomes with six previously sequenced, nearly gapless, genomes from plant and animal isolates of the FSSC (11) to study the genomic diversity of the complex. Through the definition of *Starship* mobile genetic elements and prediction of long non-coding RNAs, we uncover dramatic diversity in the FSSC and generate an important foundation for future work unravelling the dual-kingdom fungal virulence of the species complex.

## Results

### Genome sequencing of novel clinical *Fusarium solani* species complex isolates and description of the study dataset

In this study we used PacBio long read sequencing combined with Illumina short read sequencing to assemble the genomes of six novel FSSC isolates that were collected from keratitis cases by the German National Reference Center for Invasive Fungal Infections (n=3 *F. keratoplasticum* and n=3 *F. petroliphilum*). To facilitate more in-depth comparisons within the species complex, and to include isolates of environmental and animal origin, six additional FSSC isolates with similarly contiguous genomes were incorporated into the study dataset (11). The published samples included isolates of *F. falciforme*, *F. keratoplasticum, F. solani* FSSC12, *F. vanettenii*, and originated from plant (pea plant and orchid flower) and marine animal (including hawksbill sea turtle, roughtail stingray, and harbor seal) sources (Table 1). The final assemblies of the novel and published assemblies were highly contiguous, with an average of 58 scaffolds (range 14 to 143) and an average N50 of 3.8 Mb ± 0.4 (Table 1). The FSSC genomes analyzed in this study ranged in size from 48.0 Mb for one of the newly sequenced *F. keratoplasticum* isolates to 72.9 Mb for *F. vanettenii* Fs6 and encoded between 14,050 and 16,755 protein coding genes. Parallelling the range of genome sizes, a similar pattern was observed for the fraction of the genome consisting of repetitive regions, including transposable elements, which varied from 5% for *F. keratoplasticum* NRZ-2015-035 to 14% for *F. vanettenii* Fs6. Using BUSCO as a measure of genome completeness, we found that all genomes were highly complete, recovering on average >98.5% of the expected single copy orthologs. Despite originating from different studies, there were no significant differences in key genome quality metrics such as N50, the number of protein coding genes, or BUSCO completeness between the isolates sequenced in this study and the public assemblies (p>0.05 by Mann-Whitney U test).

**Table 1:** Genomic characteristics of FSSC isolates included in this study. BUSCO completeness quantified as the percent of BUSCOs identified as single copies using the hypocreales_odb10.2019-11-20 ortholog set. lncRNAs were only predicted for *F. keratoplasticum* and *F. petroliphilum* based on the availability of RNAseq data for these species.

|  | Scaffolds (n) | N50 (Mb) | Genome Size (Mb) | Protein coding genes | lncRNAs | BUSCO completeness | Repeat content | Isolation source | Reference |
| --- | --- | --- | --- | --- | --- | --- | --- | --- | --- |
| *Fusarium falciforme* Fu3 | 14 | 3.88 | 53.6 | 15,051 | - | 99.1% | 10.1% | Hawksbill sea turtle | [11] |
| *Fusarium keratoplasticum* Fu6 | 25 | 4.21 | 49.5 | 14,050 | - | 99.0% | 7.1% | Hawksbill sea turtle | [11] |
| *Fusarium keratoplasticum* LHS11 | 26 | 3.50 | 53.5 | 15,777 | - | 98.9% | 7.2% | Harbor seal | [11] |
| *Fusarium keratoplasticum* NRZ-2014-069 | 70 | 4.16 | 49.6 | 14,398 | 2,632 | 98.9% | 4.8% | Keratitis (human eye) | This study |
| *Fusarium keratoplasticum* NRZ-2015-087 | 53 | 4.02 | 50.6 | 14,387 | 2,694 | 98.9% | 7.1% | Keratitis (human eye) | This study |
| *Fusarium keratoplasticum* NRZ-2016-116 | 55 | 4.24 | 50.5 | 14,979 | 2,654 | 98.9% | 6.5% | Keratitis (human eye) | This study |
| *Fusarium petroliphilum* NRZ-2014-013 | 120 | 3.98 | 51.2 | 15,060 | 2,327 | 99.1% | 6.7% | Keratitis (human eye) | This study |
| *Fusarium petroliphilum* NRZ-2015-035 | 143 | 3.34 | 48.0 | 14,212 | 2,322 | 98.5% | 4.5% | Keratitis (human eye) | This study |
| *Fusarium petroliphilum* NRZ-2014-079 | 95 | 3.33 | 48.5 | 14,541 | 2,222 | 99.0% | 4.8% | Keratitis (human eye) | This study |
| *Fusarium* sp. FSSC12 LHS14 | 35 | 3.82 | 56.3 | 14,512 | - | 99.1% | 11.2% | Roughtail stingray | [11] |
| *Fusarium* sp. Ph1 | 24 | 3.30 | 51.5 | 14,109 | - | 99.0% | 8.6% | Orchid flower | [11] |
| *Fusarium vanettenii* Fs6 | 39 | 3.36 | 72.9 | 16,755 | - | 99.1% | 14.2% | Pea plant | [11] |

### Human and plant association is polyphyletic in the FSSC

To determine the phylogenetic relationship of the newly sequenced clinical isolates within the genus *Fusarium* and the FSSC complex, we constructed a whole genome phylogeny from amino acid sequence of conserved single copy orthologs. As expected, the new isolates of *F. keratoplasticum* and *F. petroliphilum* were located within clade 3 of the FSSC in the phylogeny (Figure 1). The *F. petroliphilum* isolates formed a monophyletic group, with *Fusarium* FSSC12 being the closest relative in the dataset. The *F. keratoplasticum* isolates belonged to a separate monophyletic group containing *F. solani sensu stricto*, *F. falciforme*, and the two other *F. keratoplasticum* isolates in the dataset. Interestingly, the clinical *F. keratoplasticum* isolates from Germany did not segregate from the marine host-associated isolates from east Asia, indicating a lack of geographic population structure or host specificity in the species. This overall pattern was observed in the entire complex, with virulence in both plant and human hosts being polyphyletic in the FSSC.

**Figure 1.**
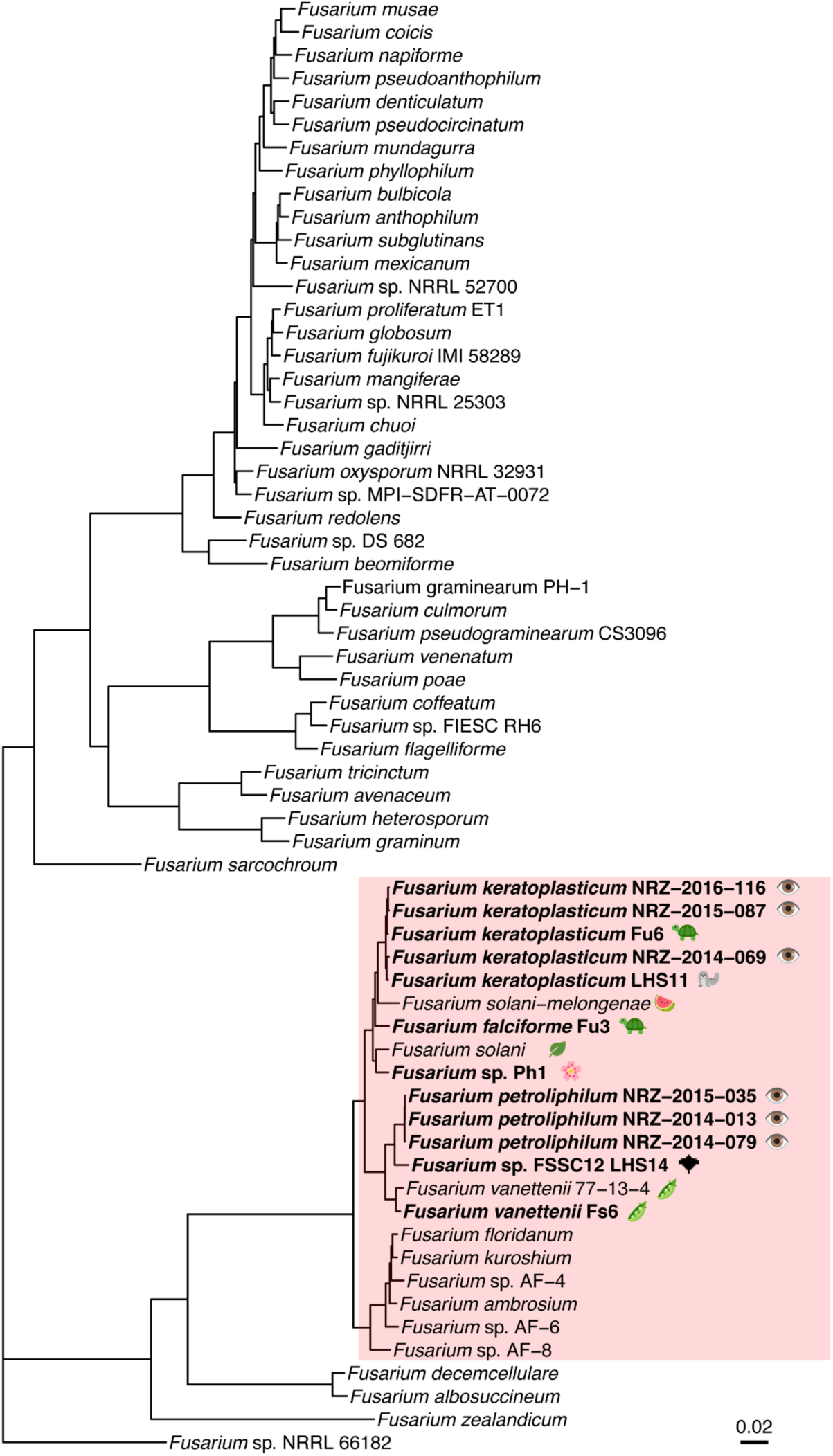
Phylogenetic reconstruction of *Fusarium* spp., including the FSSC isolates analyzed in this study. Phylogeny constructed from an amino acid sequence of 1,866 single copy orthologs. Pink box indicates the *Fusarium solani* species complex. The strains with highly contiguous long-read genome assemblies included in this study are indicated in bold. Emoji indicates the source for the FSSC isolates that belong to the same clade as the study dataset members. All branches supported by ultrafast bootstrap values >98%.

### 41% of genes are common to all FSSC isolates, while pan-gene conservation is higher in individual species

We next sought to quantify the genomic diversity of the FSSC by characterizing its pan-genome and delineating the genes common to all genomes analyzed (core genome) and those which show presence-absence variation within the species complex (accessory genome). Of the 27,068 orthologous genes identified across the 12 isolates and 6 species, only 41% (n=11,079) were core, or present in all isolates (Figure 2a, b). The FSSC pan-genome exhibited a long tail distribution, where 6% of the pan-genome (n=1,666) was present in all but one sample and 7% (n=1,946) of the accessory genome was isolate-specific (7%; n=1,946) (Figure 2b). Despite the low fraction of core genes, Heaps’ law indicates that the FSSC pan-genome is closed (alpha = 1.57, where alpha values > 1 indicate a closed pan-genome) and the number of pan-genes begins to plateau with the analysis of six genomes (Figure 2c). Protein sequences of core genes were significantly longer than those of accessory genes, averaging 505 amino acids compared to 377 for the latter (Wilcoxon p < 0.0001). Within the scope of single species, the fraction of core genes was notably higher. For *F. keratoplasticum,* 75% of pan-genes were core, or in all 5 isolates (13,275/17,617), while 88% of pan-genes were core in *F. petroliphilum* (14,067/15,923 pan-genes; n=3 isolates) (Figure 2d, e).

**Figure 2.**
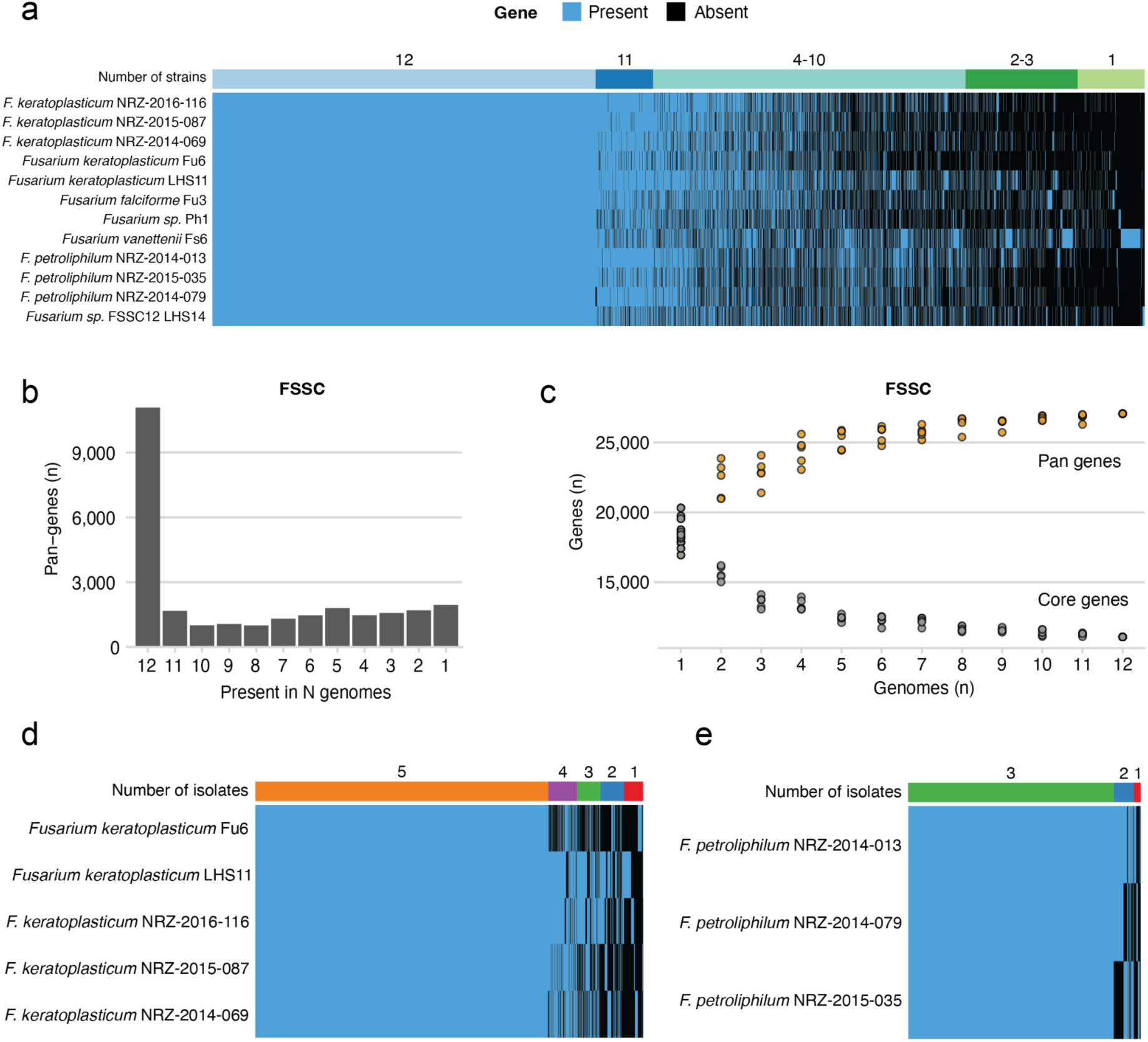
The pan-genome of the FSSC, *Fusarium keratoplasticum*, and *F. petroliphilum*. (a, d, e). Pan-gene presence-absence matrix of FSSC (a), *F. keratoplasticum* (d), and *F. petroliphilum*, where gene presence is indicated with blue and gene absence in black. Colored bars at the top of the matrices indicate the number of isolates each pangene was present in. (b) Histogram of pangene frequency in the FSSC, calculated as the number of genomes in which each pan-gene is present. (c) Pan-gene accumulation plot quantifying the relationship between the number of genomes analyzed and the number of pan-genes identified (orange) and the fraction of which were core (grey). Dots represent the individual values from 24 permutations.

### Diverging patterns in sequence conservation and repetitive elements between core and accessory chromosomes

In addition to also possessing a high degree of gene presence-absence variation, some of the accessory gene content of the *F. oxysporum* species complex is known to be encoded on discrete accessory chromosomes. These accessory chromosomes are involved in plant virulence and the ability to infect specific host plants, but their role in human infection is unclear. Intriguingly, work in the *F. oxysporum* species complex hints that clinical isolates of this species complex may carry accessory chromosomes that are distinct from plant-pathogenic isolates (19). To investigate this in the FSSC, we analyzed our study dataset for the conservation of both core and accessory contigs or chromosomes among the isolates from plant and animal origins. We identified 12 core chromosomes, which were present across the isolates in the dataset with a high degree of nucleotide identity and synteny (Figure 3). However, the relative proportion of the core chromosomes compared to the total genome varied considerably within the species complex. Core chromosomes, encompassing a total length of 46.8 Mb in *F. keratoplasticum* Fu6, made up 95% of its total genome length. In comparison, core chromosomes comprised only 73% of the total genome length of *F. vanettenii* Fs6, which contained 15 additional large contigs or chromosomes with low identity to any other sequences in the dataset (Supplementary Figure 1). We also observed that all samples in the clade containing *F. petroliphilum* and *F. sp.* FSSC12 possessed a substantial chromosomal translocation between the two largest core chromosomes (Supplementary Figure 3).

**Figure 3.**
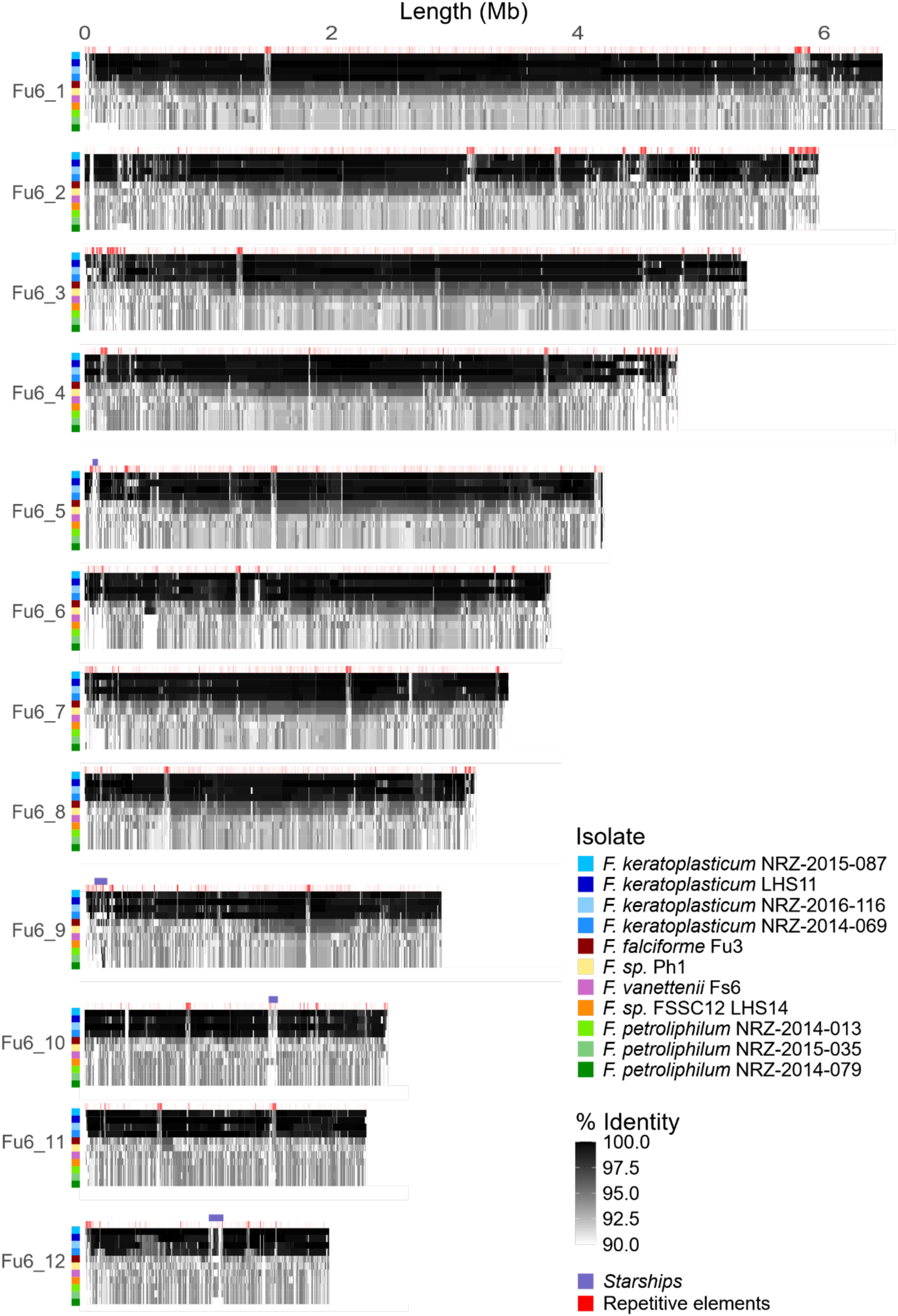
Sequence conservation of core chromosomes of the FSSC. Gray-scale fill indicates the nucleotide percent identity between the core chromosome sequence of *F. keratoplasticum* Fu6 and other FSSC members, which are indicated with the colored boxes. Violet bars on Fu6_5, Fu6_9, Fu6_10, and Fu6_12 indicate the location of *Starship* elements. Red bars indicate presence of transposable and other canonical repetitive elements.

In contrast to the high synteny and sequence conservation among the core chromosomes, the accessory chromosomes were largely species- or isolate-specific or had much lower levels of sequence identity for the regions that were present (Figure 4). We detected between three and eight accessory chromosomes or contigs >150kb in the *F. keratoplasticum* isolates, and between five and nine in the three *F. petroliphilum* isolates. Many accessory chromosomes were conserved in all isolates of the same species, but not across species. For example, two of the three accessory chromosomes present in *F. keratoplasticum* Fu6 had high identity with the other *F. keratoplasticum* isolates but low synteny with other FSSC isolates (Figure 4). In other cases, accessory chromosomes were isolate-specific, such as chromosome 13 of *F. keratoplasticum* LHS11, which displayed <52% identity with any other contig in the dataset. However, we did not identify any accessory chromosomes or contigs in *F. keratoplasticum* that were specific to the clinical strains. Notably, we also observed cases where the distribution of accessory chromosomes or contigs did not mirror the core genome-generated phylogeny, suggesting that horizontal transfer is possible in the FSSC, as has been demonstrated for *Fusarium oxysporum*. Four *F. keratoplasticum* isolates (Fu6, LHS11, 2015-087, and 2016-116) possessed one to three contigs with higher nucleotide identity to accessory genomic regions of *F. petroliphilum* than to the other strains of its own species (Figure 4a). Similarly, *F. petroliphilum* strains 2014-013 and 2015-035 possessed accessory contigs with higher synteny and nucleotide identity to *F. keratoplasticum* than to their own species (Figure 4b).

**Figure 4.**
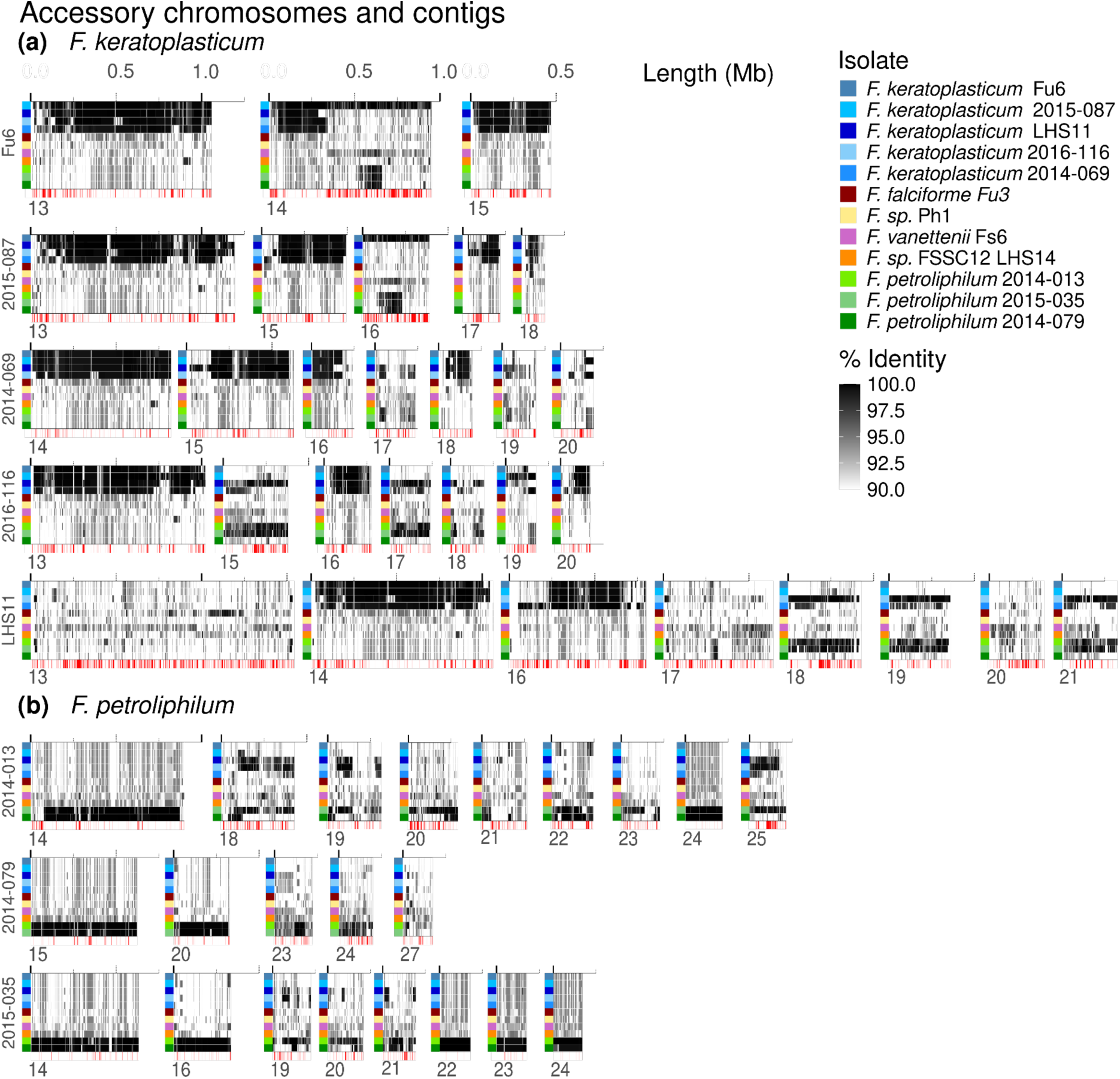
Conservation of accessory chromosomes and contigs from *Fusarium keratoplasticum* and *Fusarium petroliphilum*. (a-b) Accessory chromosomes and contigs of (a) *F. keratoplasticum* or (b) *F. petroliphilum* strains and their sequence conservation with other FSSC isolates. Shading indicates percent identity across the syntenic block between the indicated isolate and accessory chromosome/contig indicated, versus the best matching chromosome or contig from the other isolates. For simplicity, the subject (reference sequence) for each comparison is omitted from the plot as it would be a solid black band. Red lines indicate the location of transposable elements and other repetitive elements. Only genomic fragments >150 kb are displayed.

The accessory chromosomes of all six FSSC species possessed a significantly higher fraction of canonical repetitive elements, including simple repeats and class I and II transposons, compared to the core chromosomes (p <0.05 for all species by Mann-Whitney U test) (Supplementary Figure 2). As a concrete example of this, the median canonical repeat content of the 12 core chromosomes among the five *F. keratoplasticum* isolates was 7%, compared to 25% for the accessory chromosomes and contigs. Altogether, we defined 12 core chromosomes with a high degree of sequence conservation across the FSSC and an assortment of species- and isolate-specific accessory chromosomes that also had a higher content of canonical repetitive elements and the presence of which were not always congruent with the core genome phylogeny.

### 57% of intergenic lncRNAs in *F. keratoplasticum* are conserved in *F. petroliphilum*

In addition to variation in protein-coding genes, eukaryotic genomes encode myriad long non-coding transcripts (lncRNAs) that are often involved in gene regulation and evolve faster than coding genes (37). Previous work on lncRNAs in *Fusarium graminearum* has highlighted their roles in regulating sexual reproduction and carotenoid biosynthesis (38–40). However, the lncRNA landscape in *Fusarium* has not been explored beyond this. To this end, we sequenced the transcriptomes of *F. keratoplasticum* NRZ-2014-069 and *F. petroliphilum* NRZ-2014-079 conidia and of mycelia grown under several environmental conditions to capture the breadth of lncRNAs expressed by these two species. Using this approach, we predicted lncRNAs for all six of the new *F. keratoplasticum* and *F. petroliphilum* strains, identifying a range of 2,222 to 2,694 putative lncRNAs per genome (Table 1). For detailed downstream characterization, we focused on *F. keratoplasticum* NRZ-2014-069 and *F. petroliphilum* NRZ-2014-079 as representative models for each species. We found that each shared a comparable ratio of intergenic to antisense lncRNAs, with the majority of lncRNAs being intergenic and only 16% of lncRNAs being antisense in both species (Supplementary Figure 4). To identify conserved intergenic lncRNAs between the two species, we used a combination of reciprocal BLASTn (local alignment), MUMmer (global alignment), and synteny-based approaches. BLASTn identified 395 conserved lncRNA pairs, representing 18% of the intergenic lncRNAs in *F. keratoplasticum*. MUMmer alignment identified 385 pairs, with a 95% concurrence with BLASTn. As lncRNAs are known to have lower levels of sequence conservation compared to protein-coding genes, we also employed a synteny-based approach as in [34]. Conserved lncRNAs were identified between the two species if they were flanked by at least one orthologous gene both up- and downstream. This approach identified 1,647 syntenic genomic blocks, containing lncRNAs (Supplementary Figure 5, Supplementary File 3). Taking the union of the three approaches revealed that 57% of the intergenic lncRNAs in *F. keratoplasticum* (1,264/2,224) were conserved in *F. petroliphilum*, in line with the low the conservation rate among protein-coding genes between the two species. Notably, all conserved lncRNAs identified through synteny analysis were located on core chromosomes. The fact that these lncRNAs are located on core chromosomes and are conserved between species indeed suggests that they are likely to have conserved and crucial functions. However, underscoring the likely role of noncoding genes in species- and isolate differences, we also detected 51 species-specific lncRNAs in *F. keratoplasticum* and 104 in *F. petroliphilum* that were located on accessory chromosomes. Altogether, we defined the lncRNAome of two FSSC species and found that noncoding genes follow the same trend as the rest of the genome, with a low degree of conservation. However, the existence of conserved lncRNAs, despite the rapid evolution of non-coding genes, suggests that they likely perform critical functions.

### *Fusarium keratoplasticum* genomes are enriched in branched-chain amino acid and metallopeptidases, while all FSSC genomes show similar numbers of secreted effectors

Given the degree of variation in the pan-genome of the FSSC and the accessory chromosomes within and between species, we next performed gene set enrichment analysis to identify Gene Ontology (GO) categories that were over- or underrepresented in species of the FSSC, with a focus on *F. keratoplasticum* and *F. petroliphilum*. The genomes of *F. keratoplasticum* had an expansion of genes involved in branched-chain amino acid GO terms relative to other FSSC genomes, a pathway which is non-conserved in metazoans, but essential to growth and virulence in several human pathogenic fungi (Supplementary Figure 6) (41, 42). We also observed an enrichment in genes annotated with protein ubiquitination and metallopeptidase activity in this species, which serve roles in fungal growth and pathogenicity, including tissue invasion and hyphae formation in other fungal species (43–45). The genomes o*f F. petroliphilum* had higher numbers of genes with endonuclease activity and glutathione metabolic processes relative to the other FSSC species.

Underscoring their evolutionary trajectory as plant pathogens, all FSSC genomes contained a similar abundance of secreted effectors and genes with GO annotations related to plant virulence. These included similar levels of genes with GO terms related to chitosanase activity, pectinesterase activity, and cellulase activity, all of which are involved in the degradation of plant cell walls (46–48). All genomes also had between 477-546 secreted effectors. The fact that these genomic features were present at similar levels in plant isolates and human and marine animal isolates suggests that the latter likely retain some degree of plant pathogenicity or, alternatively, that these effectors are also important during interaction with higher eukaryotes. Taken together, the gene functional analysis revealed that all FSSC genomes share features associated with plant virulence, while *F. keratoplasticum* and *F. petroliphilum* possessed differential expansion of genes involved in branched chain amino acid metabolism, oxidative stress protection, and DNA repair.

### Comparison of secondary metabolite biosynthetic gene clusters reveals a core set of actively transcribed secondary metabolites in the FSSC

Fungi produce an extraordinary array of secondary metabolites, which play key roles in their interactions with their environment and other organisms. They can be used for microbial communication, nutrient scavenging, and even as antimicrobial weapons (49–51). These molecules are typically encoded in biosynthetic gene clusters (BGCs), or organized units of multiple genes all involved in metabolite production. *Fusarium* spp, including FSSC, produce a diverse range of compounds, with >160 novel *Fusarium* metabolites described (52, 53). However, the range of BGC produced by FSSC isolates from human and animal origin has not been studied. Across the 12 genomes, we predicted 45 total BGCs and each isolate encoded a median of 36 BGCs (range 35-39). Of the BGCs identified, 25 were present across all study strains, representing the core secondary metabolome of the FSSC (Figure 5). Among the core BGCs that matched a known product were the agricultural contaminant ochratoxin A, as well as squalestatin S1, metachelin, tryptoquialanine, and lucilactaene, many of which have antimicrobial activity (54–58). The BGC encoding the antimicrobial and immunosuppressant cyclosporin C was present in the phylogenetic subclade encompassing *F. keratoplasticum, F. falciforme*, and *F. sp.* Ph1, but absent in the other FSSC clade examined. Also of interest, *F. sp.* Ph1 was the sole FSSC predicted to produce gibberellin, a BGC typically restricted to the *Fusarium fujikuroi* species complex. Gibberellin is a plant growth hormone and its production by *Fusarium fujikuroi* is considered a major factor in the development of symptoms in rice bakanae disease, which causes up to 50% crop loss (59, 60).

**Figure 5.**
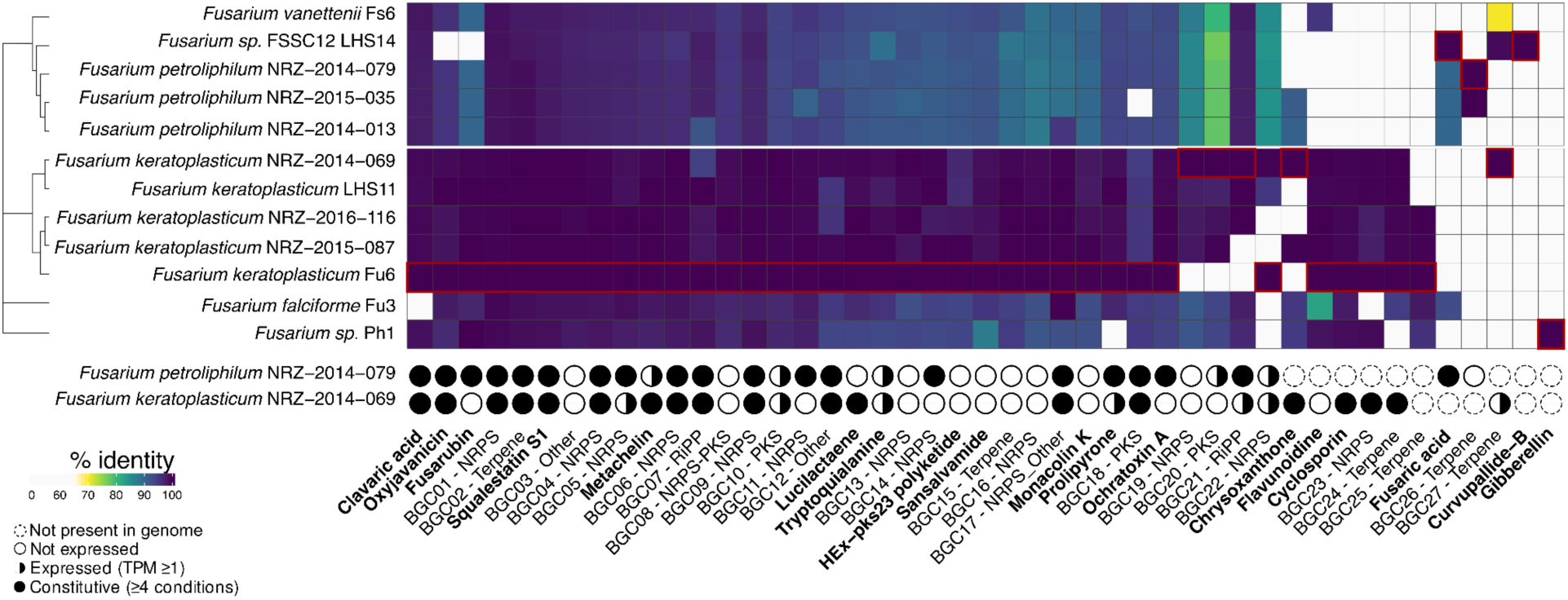
Comparison of secondary metabolite biosynthetic gene clusters reveals a core set of secondary metabolites in the FSSC and prevailing expression. BGCs with a known predicted product are indicated in bold and those with unknown products labeled with their natural product class. Percent identity was calculated using amino acid sequence of the backbone enzymes. Red boxes indicate the isolate whose sequence was used as the query when calculating percent identity. The circles beneath the heatmap indicate the breadth of conditions where transcription of the core backbone enzyme was detected in *F. petroliphilum* NRZ-2014-079 and *F. keratoplasticum* NRZ-2014-069. Dashed circles represent BGCs that were not present in the genome, empty grey circles (○) represent BGCs where no expression was detected in any of the five conditions assayed (TPM < 1), black half-filled circles (◑) indicate condition-specific gene expression with TPM ≥ 1 in 1–3 conditions, and black filled circles (●) denote constitutive expression (TPM ≥ 1 in ≥ 4 conditions). The dendrogram on the left represents the phylogenetic relationship of the isolates.

BGCs are often expressed under particular environmental stimuli or conditions. When we examined the transcriptomes of *F. keratoplasticum* NRZ-2014-069 and *F. petroliphilum* NRZ-2014-079, we observed that many BGCs were transcribed in at least one condition (n=25/40 for *F. keratoplasticum* and n=25/36 for *F. petroliphilum*; Figure 5) and many in four or five conditions. Among the ubiquitous expressed BGCs across both species were clavaric acid, oxyjavanicin, and squalestatin S1. Fusarubin was expressed in nearly all samples and conditions for *F. petroliphilum,* but none for *F. keratoplaticum*, despite the cluster being in the genome, indicating a divergence in regulation within the complex. Finally, the alkaloid metabolite tryptoquialanine was expressed at both high temperatures and under carbon starvation, and across both species, but not in complete media, suggesting a potential relevance for human infection. Collectively this suggests that the FSSC uses many of its secondary metabolites as a common response across many conditions, rather than just reserving them for when a specific stress or microbe is encountered. Altogether we find a wealth of secondary metabolites in the FSSC, including many that are actively transcriptionally transcribed but have unknown products, making them exciting candidates for future study.

### *F. keratoplasticum* has highest number of *Starship* mobile genetic elements in the FSSC

Given the pronounced degree of gene content variation between members of the FSSC and within the same species, we next sought to investigate factors underpinning this. *Starships* are a newly described class of gigantic transposable elements concentrated in the *Pezizomycotina* that can extend up to 700 kb. They have the ability to move genetic information within a genome, to other members of the species, or even to facilitate the horizontal transfer of large genomic “cargo” regions between evolutionary distant fungal species (61–63). *Starships* are defined by the presence of a site-specific recombinase, or “captain”, which catalyzes the movement and also allows for the differentiation into different classes and haplotypes (64). Examination of the 12 FSSC isolates revealed 23 *Starships*, all of which were on core chromosomes, and which were categorized into five distinct families representing the *Arwing*, *Enterprise*, *Galactica*, *Phoenix*, and *Prometheus Starships*. However, *Starship* presence was not equally distributed across the FSSC (Figure 6a). *F. keratoplasticum* had the largest number of *Starships* per genome with a median of 4 (range 1-6), while the median for the species complex was 1.5. This included up to four copies of the *Enterprise* class in members of *F. keratoplasticum*. We also observed that the presence of either canonical repetitive elements (*e.g.* shorter transposons, simple repeats) or a *Starship*s coincided with breakdown in synteny and reduced nucleotide identity on core chromosomes (Figure 3), highlighting their collective role in generating genomic novelty, even within the largely conserved core chromosomes. Also of note, no predicted effectors were located within *Starship* elements, indicating that in contrast to other phytopathogens (63, 65, 66), *Starships* do not appear to be a significant driver of plant virulence in the FSSC.

**Figure 6.**
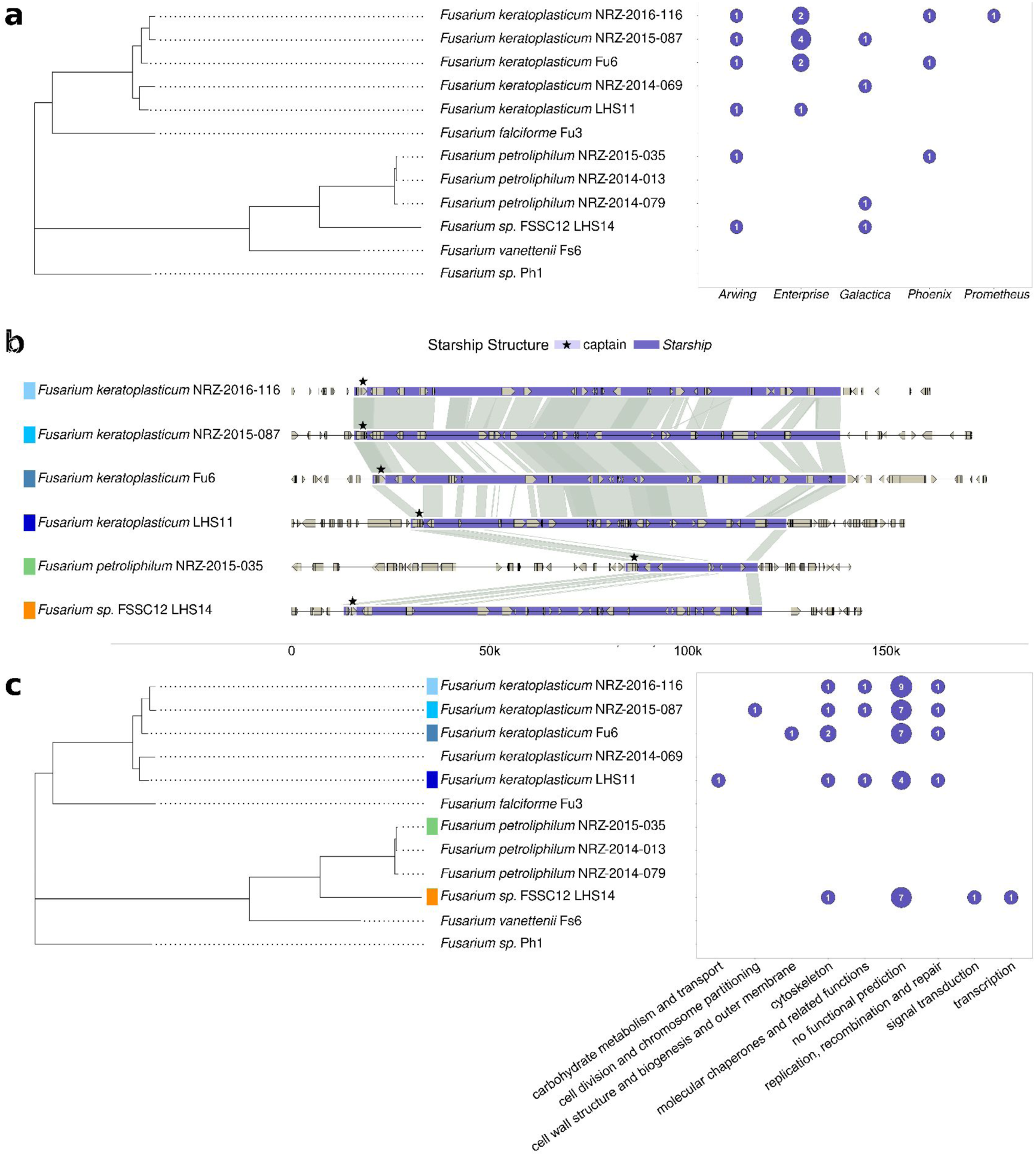
Synteny and cargo of a conserved *Arwing Starship* in the FSSC. (a) Bubble plot indicating the number of unique *Starship* elements identified per genome, categorized by class of their captain. (b) Synteny plot depicting the between the different *Arwing Starships*. The dark purple shows the whole *Starship* and the light purple in combination with a star highlights the captain of each *Starship*. Strains were ordered corresponding to their phylogenetic position in panel b. (c) Bubble plot indicating the phylogenetic relationships of the samples and the COG functional annotation of the cargo. Numbers indicate the number of ORFs with each COG annotation within the *Starship* element.

*Arwing Starships* were the most widespread across the complex and were detected as a single copy in half the samples, including some but not all *F. keratoplasticum* and *F. petroliphilum* genomes and the FSSC12. Therefore, we next focused our examination on comparing their synteny and cargo across the complex. The median *Arwing* element was 112.5 kb (range 33.4 - 122.8 kb) and contained a median of 23 genes (range: 6 - 25 genes). Synteny was highly conserved in all but *F. petroliphilum* NRZ-2015-035, which contained a truncated copy compared to the rest (Figure 6b). Notably, the presence-absence pattern of the *Arwing* element was not congruent with the genome-wide phylogeny. This indicates that either it arose in the common ancestor but has been stochastically lost by roughly half the strains or, alternatively, that it has potentially been transferred horizontally between species multiple times. Examination of the cargo functions carried by the *Arwing* element revealed genes related to the cytoskeleton, molecular chaperones, and replication, recombination and repair as the most abundant within the mobile element (Figure 6c, Supplementary Figure 7a). However, it is worth noting that the majority of the cargo genes did not contain any functional annotation, preventing a complete understanding of horizontally transferred cargo. Analysis of the *Galactica Starships* revealed the same trend in both specific functional annotation categories and the largest fraction of cargo lacking a predicted function (Supplemental Figure 7b), collectively underscoring the work still to be done in defining the functional roles of this novel class of genetic elements in the FSSC and other fungal species.

We next sought to move beyond genomic presence and assess whether *Starship*-encoded cargo was being actively transcribed. Using *F. keratoplasticum* NRZ-2014-069 and *F. petroliphilum* NRZ-2014-079 for which we had comprehensive RNA-seq data across five culture conditions, we examined gene expression in the conserved and highly syntenic *Galactica* locus shared by both strains (Supplementary Figure 6b, Supplementary Figure 8). We observed near-constitutive expression of cargo genes across most conditions in both species, with an average of 21/29 cargo genes expressed per condition for *F. keratoplasticum* and 23/25 genes for *F. petroliphilum* (Supplementary Figure 8). However, while cargo genes in *F. keratoplasticum* displayed low basal expression across all conditions assayed, *F. petroliphilum* exhibited stronger and consistent transcription across a contiguous block of hypothetical genes that also included two lncRNAs, with the strongest expression under heat stress at 34°C and may be relevant to human infection. Together, these results indicate that *Starships* have played an appreciable role in shaping the genomes of the FSSC and are not silent components of the genome, but instead exhibit condition-and species-specific transcriptional activity.

### Minimal transcriptional changes in *F. keratoplasticum* and *F. petroliphilum* at ocular infection temperatures

Given the notable genomic differences between *F. keratoplasticum* and *F. petroliphilum* and gene expression differences between a conserved *Starship* in these species, we expanded our analysis of the transcriptomes of these two species under several stressors. To this end, we compared the transcriptomes of mycelia from *F. keratoplasticum* NRZ-2014-069 and *F. petroliphilum* NRZ-2014-079 grown in complete media at 25°C, under carbon starvation at this temperature, and at elevated temperature mimicking the surface temperature of the eye (34°C). Additionally, the transcriptome of conidia was also assayed. Growth stage (conidia vs. mycelia grown on complete media) had the largest impact on gene expression with 686 and 786 differentially expressed genes (DEG) in *F. keratoplasticum* and *F. petroliphilum* respectively (Supplementary File 4, Supplementary File 5) and indicated a shared enrichment for DNA-regulated transcription and zinc ion binding in mycelia relative to conidia (Supplementary Figure 9). Relative to complete media, growing mycelia under carbon starvation resulted in a positive enrichment for aromatic amino acid metabolic genes and a negative enrichment in genes related to translation (Supplementary Figure 9; Supplementary File 4). Unexpectedly, elevated temperature (34°C) resulted in very modest change to the transcriptome, with only 27 genes differentially expressed in *F. petroliphilum* and 47 in *F. keratoplasticum* (Supplementary File 4). This suggests that both species are more than capable of growing at temperatures experienced during mammalian association or infection with minimal transcriptional alteration compared to growth at 25°C.

Consistent with an earlier study with *Fusarium graminearum* (38), both intergenic and antisense lncRNAs identified in this study had significantly lower expression relative to protein-coding genes (Mann-Whitney U test with Bonferroni correction, p < 0.0001 for both lncRNA classes in *F. keratoplasticum* and *F. petroliphilum*; Supplementary Figure 4b). However, we did identify 117 lncRNAs in *F. keratoplasticum* and 106 in *F. petroliphilum* that were uniquely differentially expressed in conidia relative to mycelia grown in complete media, suggesting they may play a role in asexual development (Supplementary File 4). Overall, the transcriptional changes of *F. keratoplasticum* and *F. petroliphilum* were similar across species, and more notably, changes during elevated temperature were minimal, indicating that human infection temperature is not a significant stressor.

### lncRNAs are potential novel regulators of secondary metabolism in *F. keratoplasticum* and F. petroliphilum

Finally, we aimed to delineate functional roles for the newly defined lncRNAs. Although lncRNAs represent a small fraction relative to the number of protein coding genes, a similar proportion of each gene type was expressed, demonstrating the broad participation of lncRNAs in cellular processes (Figure 7a). Roughly 60% of both annotated lncRNAs and protein coding genes were expressed in at least one of the five conditions assayed in both *F. keratoplasticum* and *F. petroliphilum*, the species for which RNAseq data was available. To assess the integration of lncRNAs into the gene expression network, we used weighted co-expression analysis. The global weighted degree and intramodular connectivity, measures of overall gene connectivity and connectivity within their assigned modules, respectively, was similar between lncRNAs and protein coding genes in both *F. keratoplasticum* or *F. petroliphilum*, indicating that lncRNAs are equivalently integrated into the co-expression network as protein-coding genes in both species (Figure 7b, c).

**Figure 7.**
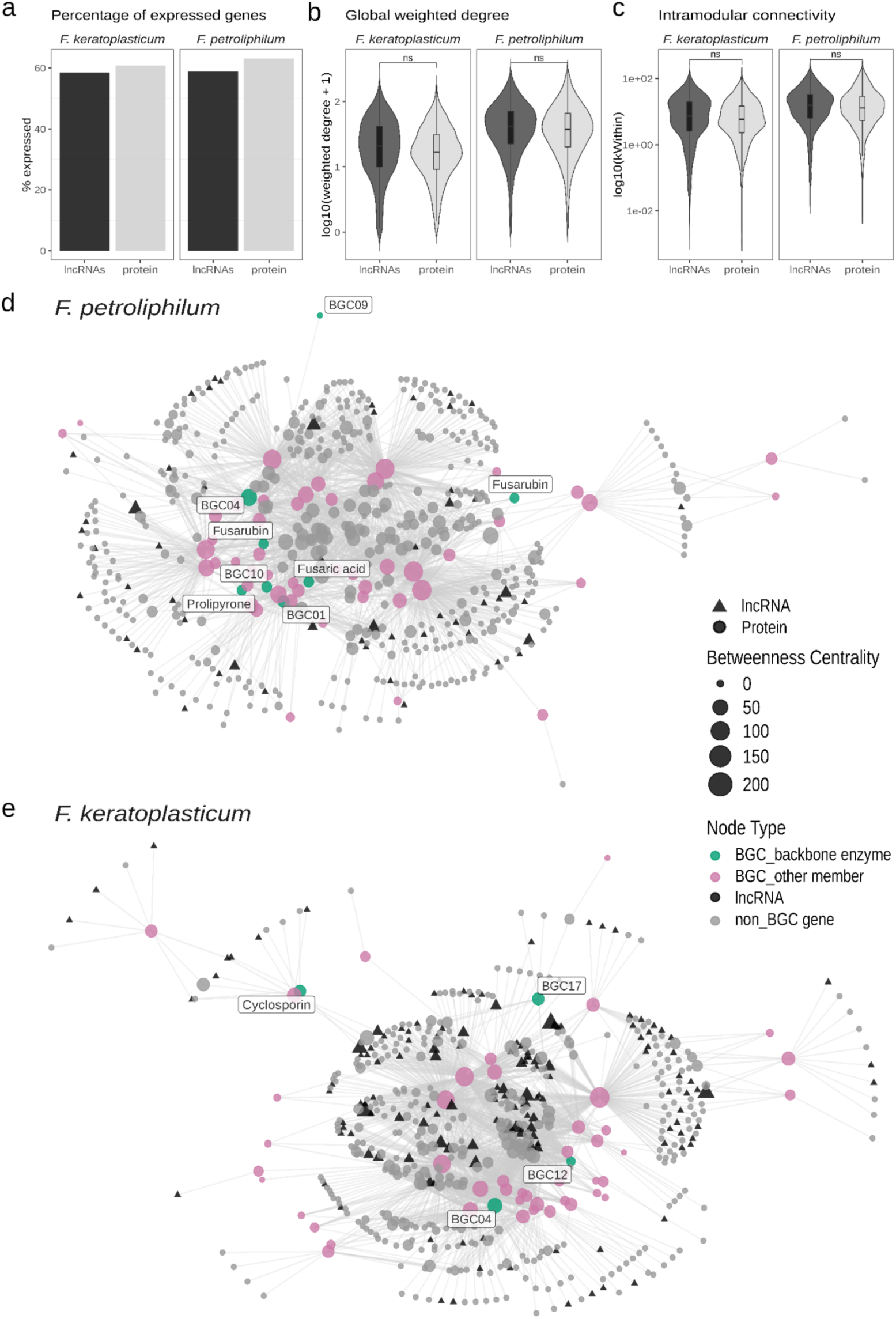
Expression and co-expression network properties of lncRNAs and protein-coding genes in *Fusarium keratoplasticum* NRZ-2014-069 and *F. petroliphilum* NRZ-2014-079. (a) Proportion of expressed genes by gene type across samples. Genes were retained for analysis if they reached a raw count threshold of ≥10 in at least 8/10 samples and their expression had a variance-stabilizing transformed (VST) value > 1 in at least one sample. (b-c) Violin plot of global connectivity (weighted degree) (b), and intramodular connectivity (kWithin) (c) within the co-expression network. Boxplot within the violin indicates the summary statistics for the distribution. (d-e) Giant connected components of filtered co-expression networks from *F. petroliphilum* NRZ-2014-079 (d) and *F. keratoplasticum* NRZ-2014-069 (e) after retaining the strongest co-expression edges. In the plot, node size is scaled by betweenness centrality and nodes are colored by gene type (BGC backbone enzyme, other BGC member, lncRNA, and non-BGC protein coding genes). Edges represent the top 0.4% (*F. keratoplasticum*) and 0.2% (*F. petroliphilum*) of TOM-based co-expression values. Statistical comparisons were performed using Mann-Whitney U test. NS indicates p > 0.05.

To identify novel pathways and regulatory functions of lncRNAs, we next examined their co-expression with protein-coding genes of known function under the well-established property that genes that are co-expressed participate in the same function or are members of the same pathway (67, 68). We specifically focused on characterizing the role of lncRNAs as contributors to secondary metabolism. The *F. petroliphilum* and *F. keratoplasticum* global co-expression networks were progressively filtered to retain only the top 0.2-0.4% of strongest co-expression edges, respectively. Within the filtered co-expression network, 29 lncRNAs in *F. keratoplasticum* and 10 lncRNAs in *F. petroliphilum* were directly connected to at least one BGC backbone enzyme, the core catalytic gene of a secondary metabolite cluster. To assess whether these lncRNA-BGC edges represent meaningful regulatory connections, we compared their co-expression strengths to those of protein-BGC interactions. Protein-BGC pairs exhibited similar median co-expression strengths as lncRNA-BGC pairs (median topological overlap measure, *ie* TOM: 0.18 vs 0.17 in *F. keratoplasticum*, 0.22 vs 0.21 in *F. petroliphilum*; Mann-Whitney U test, p = 0.11 and p = 0.71, respectively). This supports that the lncRNAs identified in our network are not simply passive associations, but interact with BGC backbone enzymes at magnitudes comparable to protein-coding genes and reinforcing that they represent a structured layer of secondary metabolism control.

The filtered co-expression networks for *F. keratoplasticum* and *F. petroliphilum* were further restricted to the giant connected component, defined as the largest subnetwork of strongly co-expressed genes (Figure 7d, e). This step reduced network fragmentation and ensured that lncRNA–BGC associations were evaluated within a single, highly connected network context. In *F. keratoplasticum*, lncRNAs connected to the cyclosporin biosynthetic gene cluster, two nonribosomal peptides or NRPs (BGC04 and BGC17), and to a phosphonate-like cluster (BGC12), suggesting roles in the regulation of these clusters. In *F. petroliphilum*, lncRNAs were directly connected with the BGCs responsible for choline, fusaric acid, fusarubin, and prolipyrone biosynthesis, as well as several BGCs with unknown products (NRPSs BGC04 and BGC09; polyketide synthase type I BGC10) (Figure 7d, e; Supplementary Table 8). Most lncRNAs were connected to a single backbone enzyme. However, a small subset displayed connectivity to two distinct backbone enzymes, suggesting they may play more global roles in regulating secondary metabolism. These included four lncRNAs in *F. keratoplasticum* (associated with BGC04 and BGC12) and two lncRNAs in *F. petroliphilum* (associated with BGC01 and BGC10) (Supplementary Table 8). Several of these multi-BGC-interacting lncRNAs were conserved across species, while others were species-specific, indicating a combination of conserved and lineage-specific regulation. Finally, we observed that while *F. keratoplasticum* and *F. petroliphilum* both contain the fusarubin BGC, it was expressed in all four mycelial conditions in the latter, but silent in all conditions in the former (Figure 5). In searching for a potential explanation, we identified a conserved lncRNA that was present in both species but whose expression was negatively correlated with the expression of the fusarubin backbone enzyme (Pearson R^2^ = 0.71, p < 0.001). We hypothesize that this lncRNA may act as a negative regulator of fusarubin biosynthesis, highlighting the value of our approach for identifying candidates for functional testing in follow-up work. Together, these results suggest that lncRNAs represent a well-integrated layer of regulation of secondary metabolism in the FSSC, with most regulating a single BGC, and a limited subset acting as more global regulators across multiple BGCs.

## Discussion

In this study we generate highly contiguous genomes for six FSSC isolates of clinical origin, including the first three genomes available for *F. petroliphilum*. We show that German keratitis isolates of *F. keratoplasticum* are not phylogenetically distinct from Asian isolates from marine animal origins. We also reveal a low level of core gene conservation among isolates of the FSSC, including pronounced variation in lncRNAs, secondary metabolites, and the presence of accessory chromosomes with isolate- and species-specific genes and a higher repeat content. Finally, we define *Starship* elements across the FSSC, finding that they are widespread, particularly in *F. keratoplasticum*, and that their cargo is actively transcribed. Finally, we define the lncRNA repertoire of the FSSC and find that these non-coding genes are broadly integrated into gene expression networks to a similar degree as protein-coding genes, including that of secondary metabolism, and likely play important, but yet to be characterized, cellular roles.

In line with previous work, our study highlights the considerable genomic flexibility of *Fusarium* spp. However, we find that the level of gene conservation within the FSSC (41%) is even lower than the *Fusarium oxysporum* species complex (FOSC), whose core genome fraction has been reported at 51-53% (69, 70). The proportion of core genes in the FSSC much also lower than environmental human fungal pathogens such as *Aspergillus fumigatus* (69%) and *Cryptococcus neoformans* (80%), but similar to another multi-kingdom pathogen, *Aspergillus flavus* (41-58%) (71–74). This may indicate that the FSSC and other multi-kingdom pathogens occupy a broader niche space and the larger pan-genome has evolved to cope with a more variable set of environmental conditions, as has been found in bacteria (75). It is also tempting to hypothesize that this expanded genomic repertoire, and the resulting flexibility underpin the ability of these organisms to infect multiple kingdoms, with very different metabolic environments and immune systems; a point that requires further investigation. On the other hand, comparing pan-genomes across studies is not without its own limitations (76, 77). It is complicated by differences in the number of isolates between studies, by pangenome methodology (*e.g.* whether consistent gene prediction methods were used, how orthologous genes are identified and what fraction of isolates must contain a gene for it to be considered ‘core’), and whether the study defines taxonomic groups using *forma specialis* or species complex definitions.

Previous work comparing a phytopathogenic *F. oxysporum* strain with two *F. oxysporum* isolates from human disease suggested that the latter may have unique accessory chromosomes. While all FSSC genomes we examined had accessory chromosomes with higher repeat content, as has been observed for other *Fusarium* species complexes (11, 18, 78, 79), we did not find clear evidence that accessory chromosome content differed from that of plant or other non-clinical sources. While we did detect accessory chromosomes that were specific to the clinical isolates of F. *petroliphilum*, our dataset lacked a phylogenetically matched comparator from environmental origin for this species. As sequencing becomes even more accessible, we expect that larger datasets of isolates from plant and human disease will help to further elucidate the important question of whether there are specific FSSC genomic elements associated with human infection.

*Starships* have facilitated the evolution of virulence in multiple phytopathogenic fungi. In the *Fusarium oxysporum* species complex, a *Starship* was responsible for the horizontal transfer of effector genes conferring host susceptibility across species boundaries (66). *Verticillium dahliae* displays evidence of anciently acquired virulence genes, whose transfer was mediated by *Starship* elements from *Fusarium* spp. (65). The wheat virulence factor ToxTA has also been transferred between multiple plant pathogens (63). In contrast, we did not find any predicted effectors or genes previously linked to FSSC pathogenicity within *Starship* elements. This suggests that while *Starships* have shaped their genome evolution, they do not seem to be the driver of virulence that they have been in other fungal pathogens. However, due to their large size, highly contiguous genomes are needed for detecting *Starships*, which are still limited in the FSSC, so this could change with the availability of additional long read genomes.

lncRNAs are increasingly recognized for their important roles in regulating pathogenicity and virulence in fungal pathogens of plants and humans, acting largely as transcriptional regulators (80–83). The list of lncRNAs described in this study represent an important starting point for determining their roles in regulating basic cellular processes in the FSSC and potentially plant and human virulence. We have already started to address this through the identification of several lncRNAs that act as putative regulators of secondary metabolism genes. In fact, as lncRNAs are not bound by the amino acid triplet code, they evolve faster than coding genes, and we posit that they may represent an important mechanism for fungal adaptation to new hosts and to environmental change, ultimately shaping the distinct pathogenic capabilities of each species. Future experimental validation of lncRNAs will help determine the mechanism by which specific transcripts regulate secondary metabolic processes and beyond.

In closing, the FSSC has the potential to be an excellent model system for studying the molecular mechanisms of multi-kingdom pathogenicity, but is currently hampered by a lack of available resources. The strains and data provided here, including the novel description of lncRNAs and *Starship* regions and identification of conserved and species- or isolate-specific genes, are an excellent resource to start to unravel multi-kingdom virulence. Through future comparisons of the transcriptomes of these isolates during plant and mammalian infection in order to identify *in planta* virulence strategies, which are repurposed for human pathogenicity we can begin to not only define mechanisms of multi-kingdom virulence but also potential evolutionary trade-offs. This is increasingly important as climate change is expected to increase the number of environmental fungal species that can grow at human body temperatures, potentially opening the door for other phytopathogens to become human pathogens.

## Materials and methods

### Strains analyzed in this study

Six isolates originating from cases of keratitis collected by the German National Reference Centre for Invasive Fungal Infection (NRZMyk) were sequenced in this work. Prior to whole genome sequencing, species identification was performed by sequencing of the TEF-1α gene as in (8). The additional FSSC genomes analyzed in this study originate from plant and aquatic animal hosts and were generated in (11). The accession numbers of the FSSC genomes analyzed in this study and those included in the phylogeny are available in Supplementary File 1.

### Nucleic acid isolation and sequencing

For the isolates sequenced in this study, mycelia were grown shaking at 30°C in complete media. Mycelia were ground in liquid nitrogen, heat shocked for 1 minute at 95°C, and cellular debris precipitated using steps and reagents from the MasterPure Yeast DNA Purification Kit (Lucigen). DNA was finally purified using the Monarch Genomic DNA Purification Kit (New England Biolabs) for binding and elution.

### Genome assembly and annotation, including lncRNA annotation

The genomes generated in this study were assembled using the PacBio long reads and Flye v2.9.1 (84). The draft assemblies then underwent two rounds of polishing using Pilon v1.24 (85) and the strain-matched Illumina data. Assemblies were then scaffolded using RagTag v2.1.0 (86) with minimum confidence score set to 0.5, minimum location confidence score set to 0.35, and the minimum orientation confidence score set to 0.6. The *F. keratoplasticum* genomes were scaffolded using *F. keratoplasticum* LHS11 as a reference as it was the most contiguous genome available for the species. As there was no existing chromosome-level genome available for *F. petroliphilum*, their genomes were scaffolded using LHS14 because it resulted in the highest mean location and orientation confidence scores of the FSSC isolates analyzed in this study. Sequence contamination was identified and removed from genome assemblies prior to gene prediction using Foreign Contaminant Screen (FSC) v0.4.0 (87).

Before annotation, assemblies were cleaned to remove contigs shorter than 1kb and those with >95% percent identity and >95% coverage with other contigs in the assembly. A custom FSSC repeat library was generated using RepeatModeler v2.0.4 (88) and the FSSC genome assemblies analyzed in this study. RepeatMasker v4.1.2 (89) and the FSSC repeat library were then used to identify and mask repetitive regions. Genomes were annotated using Funannotate v1.8.9 and all subsequent software referenced uses the versions packaged in this containerized version. Briefly, an *ab initio* FSSC gene prediction model was trained using Augustus and the assembled transcriptome of *Fusarium sp.* FSSC11 available from Mycocosm as the RNAseq data generated in our study was not available yet. Gene prediction was then performed using EvidenceModeler incorporating predictions from Augustus, GeneMarkES, Snap, and GlimmerHMM. tRNAs were predicted using tRNAScanSE. When strain-specific RNAseq data became available later, gene models were corrected using the update function of Funannotate, which also added 5’ and 3’ UTRs to the gene predictions. Functional annotations were assigned using PFAM, Eggnog-mapper, InterProScan, CAZYmes, BUSCO (dikarya_odb9 lineage), SignalP and KEGG. EffectorP 3.0 was used to predict secreted effectors. To ensure consistent genome annotations across the study dataset, the genome assemblies from Hoh *et al.* 2022 (11) were downloaded from NCBI and annotated as above, with the Augustus model trained using the RNAseq data generated in the same publication. Genome completeness in Table 1 was determined using BUSCO v5.2.2 and the hypocreales_odb10 lineage set.

To identify lncRNAs, we used RNA-seq data from *F. keratoplasticum* NRZ-2014-069 and *F. petroliphilum* NRZ-2014-079. For strains lacking RNA-seq data (*F. keratoplasticum* NRZ-2015-087, *F. keratoplasticum* NRZ-2016-116, *F. petroliphilum* NRZ-2014-013, and *F. petroliphilum* NRZ-2015-035), lncRNAs were predicted using data from the closely related strain due to high genome similarity, allowing accurate transcript reconstruction across strains. Genome-guided transcriptome assembly was performed using Stringtie v.2.2.1 (90) and the individual transcript assemblies from each species unified into a single transcript assembly using the Stringtie - merge option. To identify novel transcripts, assembled transcripts were compared against protein-encoding transcripts using Gffcompare v0.12.6 and novel transcripts >200 bp selected. Where applicable, the longest isoform was chosen utilizing gtf2gtf from CGAT apps v0.6.0 (91). Coding potential prediction was performed using two software tools: CPC2 v3 (92), which was executed against the UniProt database (https://www.uniprot.org/download), and FEELnc v0.2 (93). Transcripts were classified as lncRNAs when identified as non-coding transcripts by both tools. lncRNA transcripts were categorized as intergenic (class code ‘u’) or antisense (class code ‘x’) based on the Gffcompare output. To evaluate the expression levels of lncRNAs in Supplementary Figure 4, read counts for lncRNA and protein-coding transcripts were calculated using Featurecounts v2.0.1 (94). Subsequently, count data was normalized by transcript length and library size, yielding transcripts per million (TPM) values.

### Whole-genome phylogeny

The *Fusarium* spp. phylogeny was inferred from 1,164 single copy orthologs identified in all species genomes using OrthoFinder v2.5.4 (95). Amino acid sequences for each orthogroup were aligned using MUSCLE v3.8 (96). The alignments trimmed were using ClipKIT v1.3.0 (97), and then concatenated into an alignment of length 614,941. The phylogeny was constructed using IQ-TREE v2.2.0.3 (98). ModelFinder within IQ-TREE was used to identify JTT+F+I+GR as the best fitting substitution model based on Bayesian information criterion (BIC). Bootstrapping was performed using 1,000 ultrafast bootstraps and all branches were supported by values of >98%.

### Pan-genome analysis and identification of core vs. accessory chromosomes and contigs

GENESPACE v1.3.1 (99), incorporating Orthofinder v2.5.4 and MCScanX v01.11.2022, was used for synteny-guided orthologous gene identification and their presence-absence between strains. Heap’s Law calculation was done using the MicroPan v2.1 R package using 500 permutations. Chromosomes and contigs were defined as core and accessory based on alignment using the nucmer tool of MUMmer v4.0.0 (100). For these analyses, *F. keratoplasticum* Fu6, was used as the reference query. Core chromosomes were identified using a cutoff of >92.5% mean identity across a minimum alignment length of 1.75 Mb in all strains and were in agreement with those identified in Hoh et al. 2022 (11). The remaining chromosomes and contigs, which were not found in all 12 study strains or which showed <92.0% identify, were designated as accessory.

To compare the intergenic lncRNAs of *F. keratoplasticum* and *F. petroliphilum*, we used NRZ-2014-069 and NRZ-2014-079 as representative examples of each species. Cross-species lncRNA conservation was assessed using three approaches: BLASTn (local alignment), MUMmer v.4.0.0 (global alignment) and orthology-based synteny. First, we generated two FASTA files containing the sequences of intergenic lncRNA in *F. keratoplasticum* and *F. petroliphilum*. The pairwise comparison of reciprocal hits using BLASTn and the dnadiff tool from the MUMmer4 suite were filtered to get those with e-value > 1e-03 and a coverage > 50%. In parallel, a synteny based orthology approach, developed and validated in Pegueroles *et al*. 2019 (101), was performed. We implemented a custom pipeline modificatied to fit the analysis for two species instead of four, and using the parameters ‘3 3 1’. This configuration involved analyzing three protein-coding genes on either side of a specific lncRNA, requiring at least three common genes for each pairwise comparison between species, and ensuring a minimum of one common gene on each side of the lncRNA. To visualize the alignment coordinates, we used Circos v.0.69 to plot the 1-to-1 coordinates (coordinates available in Supplementary File 3).

### RNA isolation, sequencing, and transcriptome analysis

Two representative keratitis samples, *F. keratoplasticum* NRZ-2014-069 and *F. petroliphilum* NRZ-2014-079 were selected for transcriptomic analysis under a set of five growth conditions: mycelia grown for 12h in complete media (5.0 g/L ammonium sulfate + 10.0 g/L glucose) at 25°C, carbon-starvation media (0.1 g/L glucose), elevated temperature (34°C), and ungerminated conidia harvested after 3 days’ growth at 28°C. RNAs from two replicates were isolated using a phenol-chloroform protocol. Briefly, harvested mycelia were pelleted and bead-beaten in a mix of lysis buffer and acid phenol:chloroform:isoamyl alcohol (PCI). Lysate was centrifuged, upper phase transferred, then centrifuged again in fresh PCI. The upper phase was incubated at −20°C in sodium acetate and 2-propanol, pelleted, then twice ethanol-washed. The washed pellet was air-dried and resuspended in RNAse-free dH_2_O, then stored at −80°C.

Illumina TruSeq stranded library preparation and sequencing was performed by the University of Würzburg Core Unit Systems Medicine on the NextSeq platform. Raw FastQ data files were trimmed via fastp v0.23.4 (102), quality threshold ≤20. Filtered reads were mapped using STAR v2.7.10b (103), counted using featureCounts v2.0.1 (94) and differentially expressed genes were identified using DESeq2 v1.44.0 (104), with DEGs that met the threshold of FDR p-adj < 0.01 and a lfcThreshold > 1.5.

For the functional analysis of differentially expressed genes, gene set enrichment analysis was conducted the R package fgsea v1.35.3 (105), and GO.db v3.19.1. fgsea was run using the fgseaMultilevel command with the seed 202507, the column “stats” given by DESeq2 as rank input, and filtering for pathways with a maximum gene size of 500. Terms with a padj < 0.1 were retained and collapsed if considered similar by the function “collapsePathways”. Terms were plotted only if padj < 0.01 in at least one condition. Terms were ordered in the dotplot visualization according to their Normalized Enrichment Score (NES) value, using the function hclust.

### Functional enrichment annotation and BGC analysis

To determine the functional differences between FSSC genomes, genes were converted into matching ontology terms and summed for each strain. ANOVA and post-hoc Dunnett test were then applied to identify annotations over- or under-represented in a single species, using a Bonferroni-correct p-value cutoff of 0.05. For visualization purposes per-species expression values were normalized to z-scores, representing the number of standard deviations above/below the mean.

Biosynthetic gene clusters were predicted for each isolate genome using the fungal version of antiSMASH v7.1.0 (106), and unknown metabolites were grouped together across isolates using BiG-SCAPE v1.1.5 (107). For each predicted BGC cluster, percent identity was calculated via BLASTp. For assessing whether BGCs were expressed, the cluster was classified as expressed if at least one biosynthetic backbone enzyme had TPM ≥ 1 across both replicates in at least one condition.

### *Starship* detection and synteny assessment

*Starships* were identified using starfish v1.0.4 (108) using the pre-computed orthogroups identified by OrthoFinder v2.5.5 above. Synteny among the *Starships* identified was calculated and visualised using minimap v2.24 (109) and the R package “gggenomes” (v1.0.1). Functional annotation of *Starship* cargo was based on COG predictions.

### Weighted co-expression network and BGC co-expression analysis

Weighted gene co-expression network analysis was performed on the normalized expression matrix using WGCNA, v1.73 (110) to identify modules of co-expressed genes. Samples were evaluated via hierarchical clustering and principal component analysis (PCA), and outlier genes were removed using the goodSamplesGenes function. Soft-thresholding powers were selected based on scale-free topology (R² ≥ 0.80), while maintaining adequate mean connectivity: β = 26 for *F. petroliphilum* NRZ-2014-079 and β = 30 for *F. keratoplasticum* NRZ-2014-069. Networks were constructed using signed adjacency and topological overlap (TOM) similarity, and modules were identified using blockwiseModules (TOMType = “signed”, mergeCutHeight = 0.25, maxBlockSize = 16500).

Biosynthetic gene clusters (BGCs) were annotated as described above and classified as backbone or other members based on antiSMASH functional assignments. Pairwise co-expression values were extracted from the topological overlap matrix (TOM) for all BGC-associated genes and their interactions with lncRNAs and other protein-coding genes present in the network. To reduce network complexity and remove weak associations, an initial pre-filter was applied by retaining only interactions above the 70th percentile of TOM values. A subnetwork was then constructed from this filtered edge set, where edges were further refined using a stringent, species-specific cutoff corresponding to the upper 0.2–0.4% of TOM values, accounting for differences in network size and correlation structure between species. Subnetworks were treated as undirected graphs, and nodes were categorized as lncRNAs, BGC backbone enzymes, other BGC members, or non-BGC protein-coding genes. Giant connected components were extracted, and node centrality was quantified using degree and betweenness centrality. Highly connected BGC backbone enzymes and their associated lncRNAs were highlighted as candidate regulatory hubs involved in secondary metabolism. Networks were visualized using ggraph v2.2.1 and tidygraph v1.3.1.

## Acknowledgements

Computational work in this study was performed on the HPC cluster of the Friedrich Schiller University.

## Data and regent availability

The strains sequenced in this study are deposited in and publicly available from the Jena Microbial Resource Collection (JMRC). The PacBio, Illumina WGS, and RNAseq data generated in this study are deposited in the NCBI SRA under BioProject PRJNA1137914 and will be made publicly available upon publication.

## Author Contributions

AEB, RM, and OK conceptualized the study. AZ, AS, and RM performed experimental investigations. PJTB, MLF, MP, LMK, and AEB performed formal analyses. PJTB, MLF, MP, LMK, and AEB were responsible for visualization and writing the original draft. All authors were involved in reviewing and editing the manuscript.

---

**Supplementary Figure 1.**
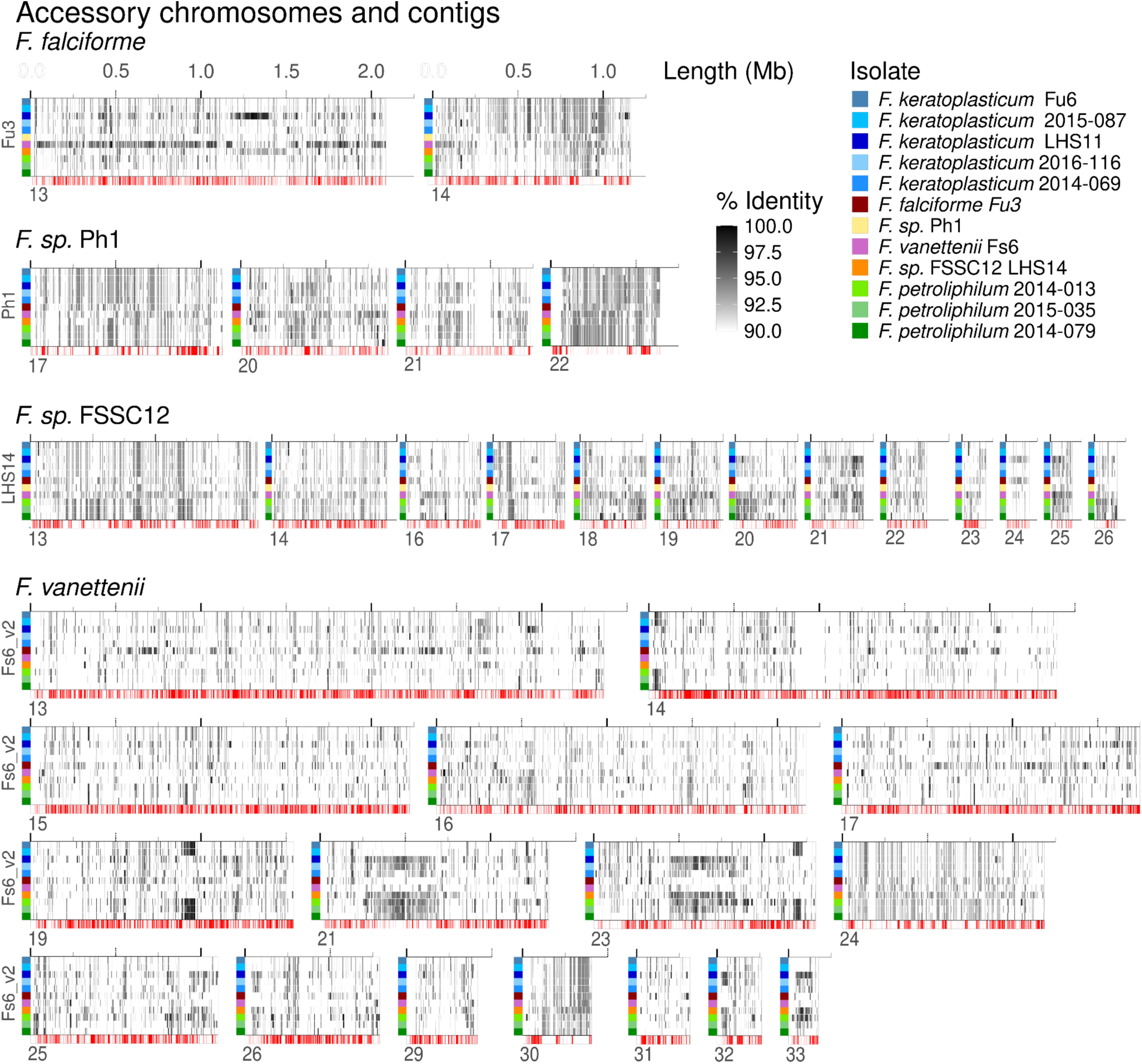
Conservation of accessory chromosomes and contigs from *Fusarium falciforme, Fusarium sp.* FSS12*, Fusarium sp.* Ph1 *and Fusarium vanettenii*. Accessory chromosomes and contigs of *F. sp.* Ph1, *F. falciforme*, *F. sp* FSS12, and *F. vanettenii* strains and their sequence conservation with other FSSC isolates. Shading indicates percent identity across the syntenic block between the indicated isolate and accessory chromosome/contig indicated, versus the best matching chromosome or contig from the other isolates. Red lines at the bottom of each subplot indicate the location of transposable elements and other repetitive elements.

**Supplementary Figure 2.**
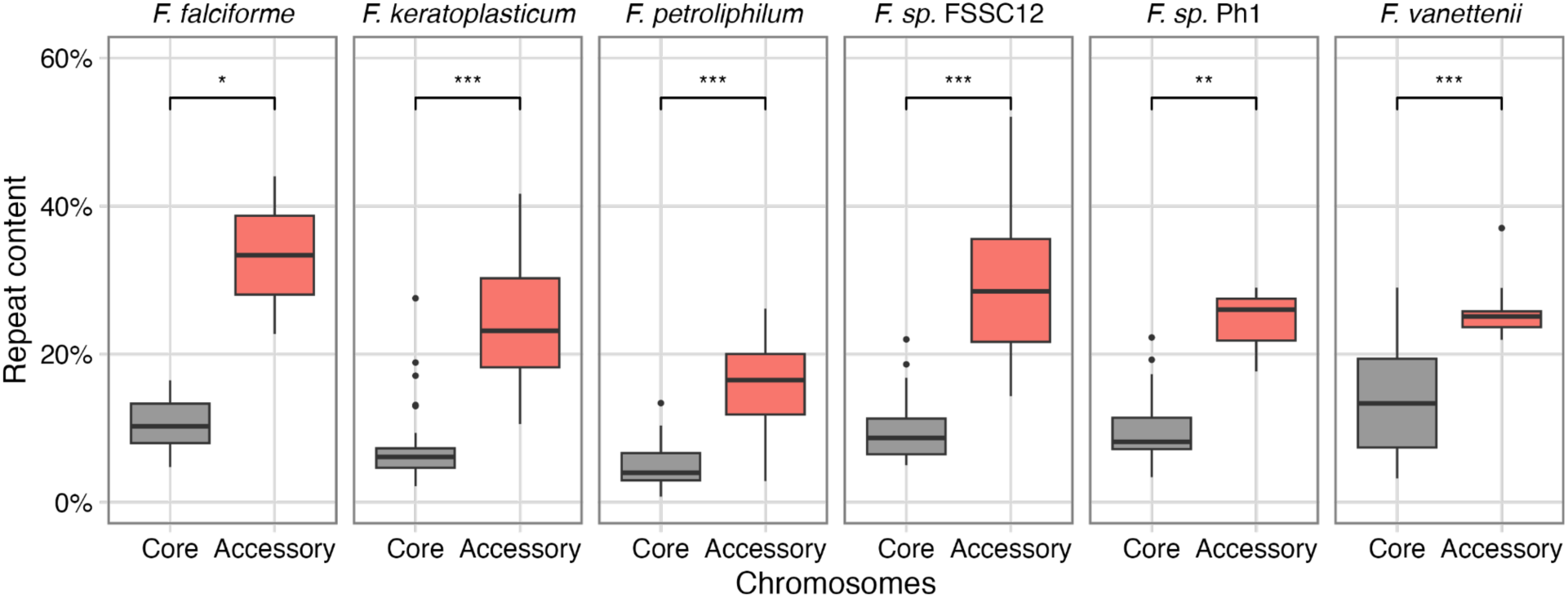
Repeat content of core and accessory chromosomes. Repeat content calculated as the number of repetitive bases masked compared to the total chromosome length. Only chromosomes and contigs >150kb were included in the analysis. *F. keratoplasticum* and *F. petroliphilum* boxplots represent aggregated chromosome data from n=3 genomes each; all other species represent the chromosomes from n=1 genome. Significance determined by Mann-Whitney U test. * denotes p<0.05, ** p<0.01, *** p<0.001.

**Supplementary Figure 3.**
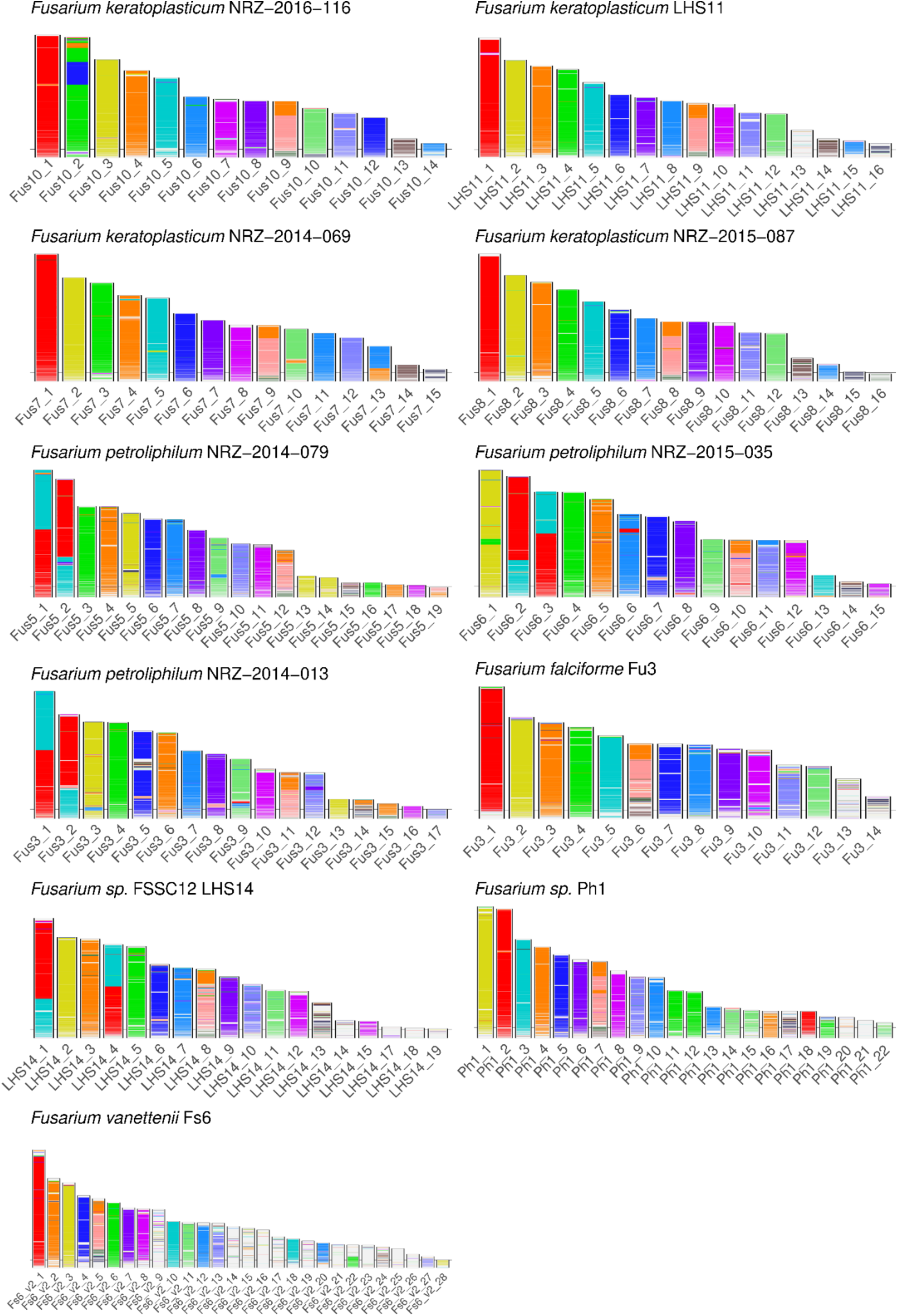
Pairwise synteny of *F. keratoplasticum* Fu6 and the other study isolates. Colors indicate synthetic blocks shared between *F. keratoplasticum* Fu6 and the indicated strain, with the individual chromosome colors representing the syntenic block’s chromosome number in Fu6. Only genomic chromosomes and contigs larger than 400 kb are displayed.

**Supplementary Figure 4.**
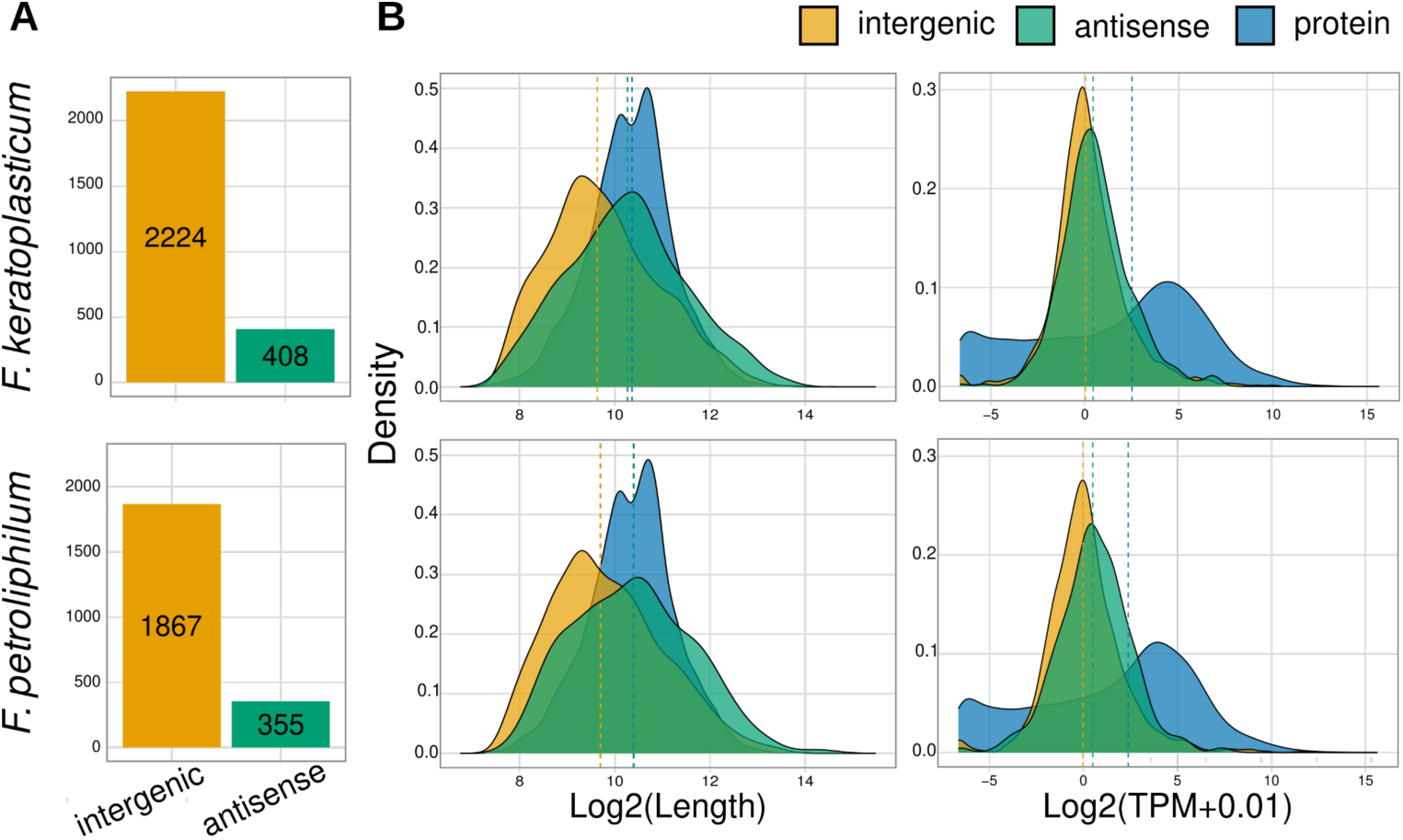
Identification and molecular properties of lncRNAs in *Fusariarium keratoplasticum and Fusarium petroliphilum*. (a) The number of intergenic and antisense predicted for *F. keratoplasticum* NRZ-2014-069 (upper panel) and *F. petroliphilum* NRZ-2014-079 (bottom panel). (b) Density plots showing the distribution of sequence length (left) and expression (right) between intergenic lncRNAs, antisense lnRNAs, and protein-coding genes. Dashed lines represent the median. Statistical significance calculated using the Mann-Whitney U test with Bonferroni correction. Intergenic lncRNAs are significantly shorter than protein-coding genes (*F. keratoplasticum*: p < 0.0001, *F. petroliphilum*: p < 0.0001). Antisense lncRNAs do not differ significantly in length compared to protein-coding genes. Both intergenic and antisense lncRNAs show lower expression levels compared to protein-coding genes (p < 0.0001 for both classes of lncRNAs in both species compared to protein coding genes.

**Supplementary Figure 5.**
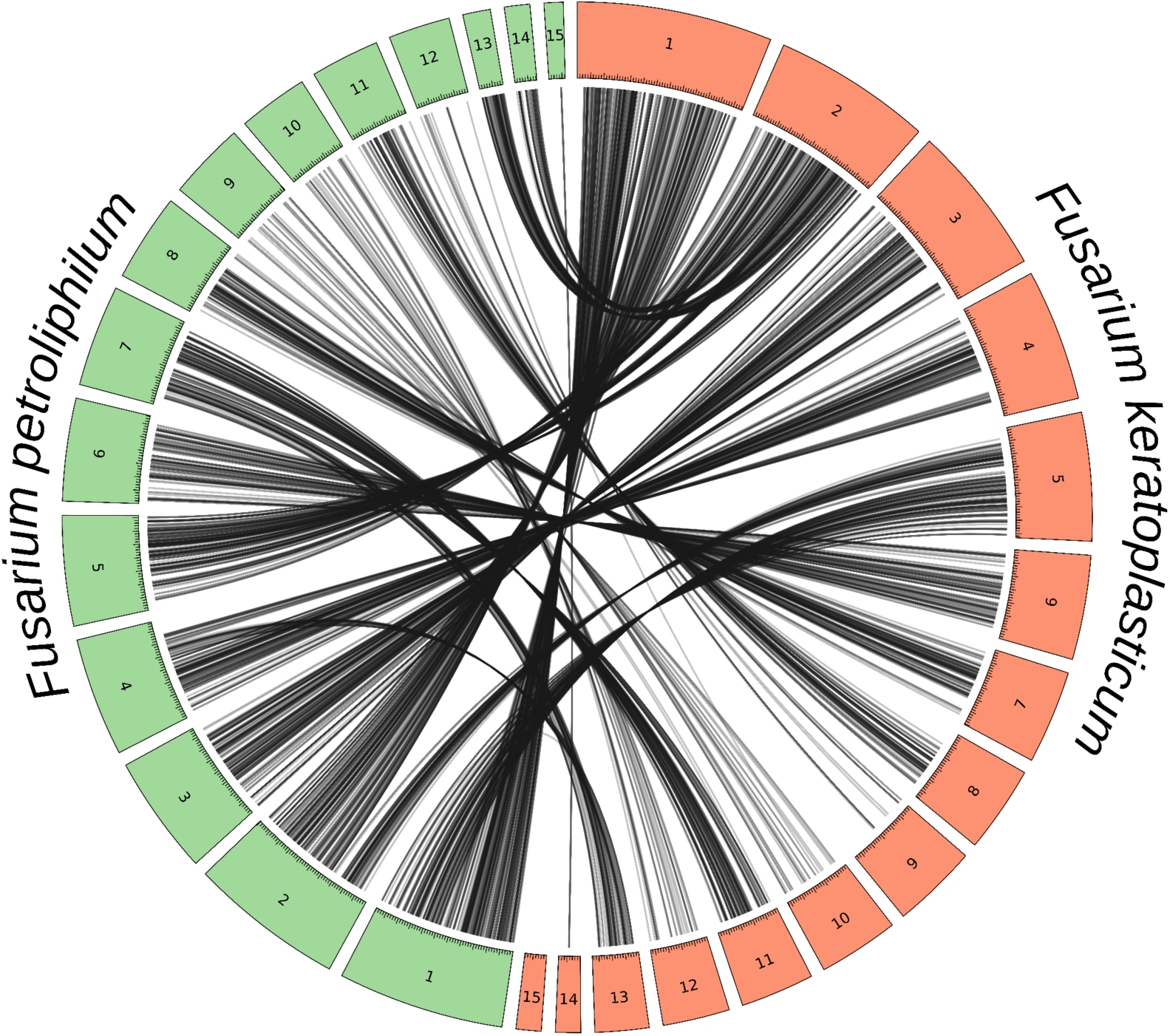
Synteny analysis of conserved intergenic lncRNAs in *Fusariarium keratoplasticum* and *Fusarium petroliphilum*. Circos plot depicts the syntenic relationships of intergenic lncRNAs between *F. keratoplasticum* NRZ-2014-069 and *F. petroliphilum* NRZ-2014-079. For a lncRNA to be considered syntenic, at least three common genes for each pairwise comparison between species, with a minimum of one common gene on each side of the lncRNA. Syntenic regions are visualized as black ribbons connecting segments between the two species. Only genomic scaffolds >615 kb are displayed.

**Supplementary Figure 6.**
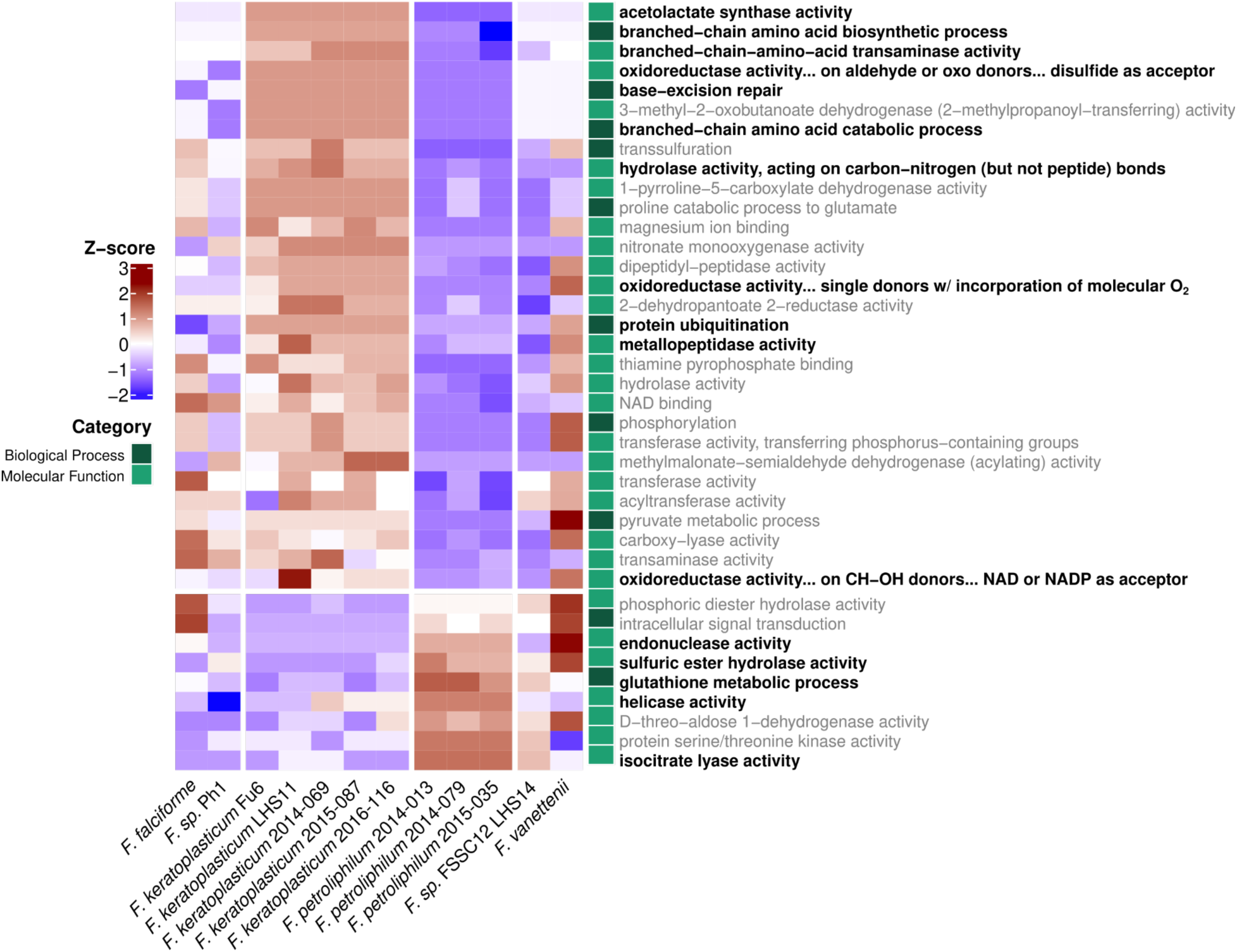
Heat map of over and underrepresented gene ontology (GO) terms in *Fusarium keratoplasticum* and *petroliphilum* compared to other members of the FSSC. Significantly over- or under-represented GO terms for *F. keratoplasticum* and *F. petroliphilum* identified by t-test and Bonferroni-corrected p < 0.05. Heat map fill represents the Z-scaled number of genes per genome annotated with each term. GO terms of particular interest or highlighted in the text are written in bold.

**Supplementary Figure 7.**
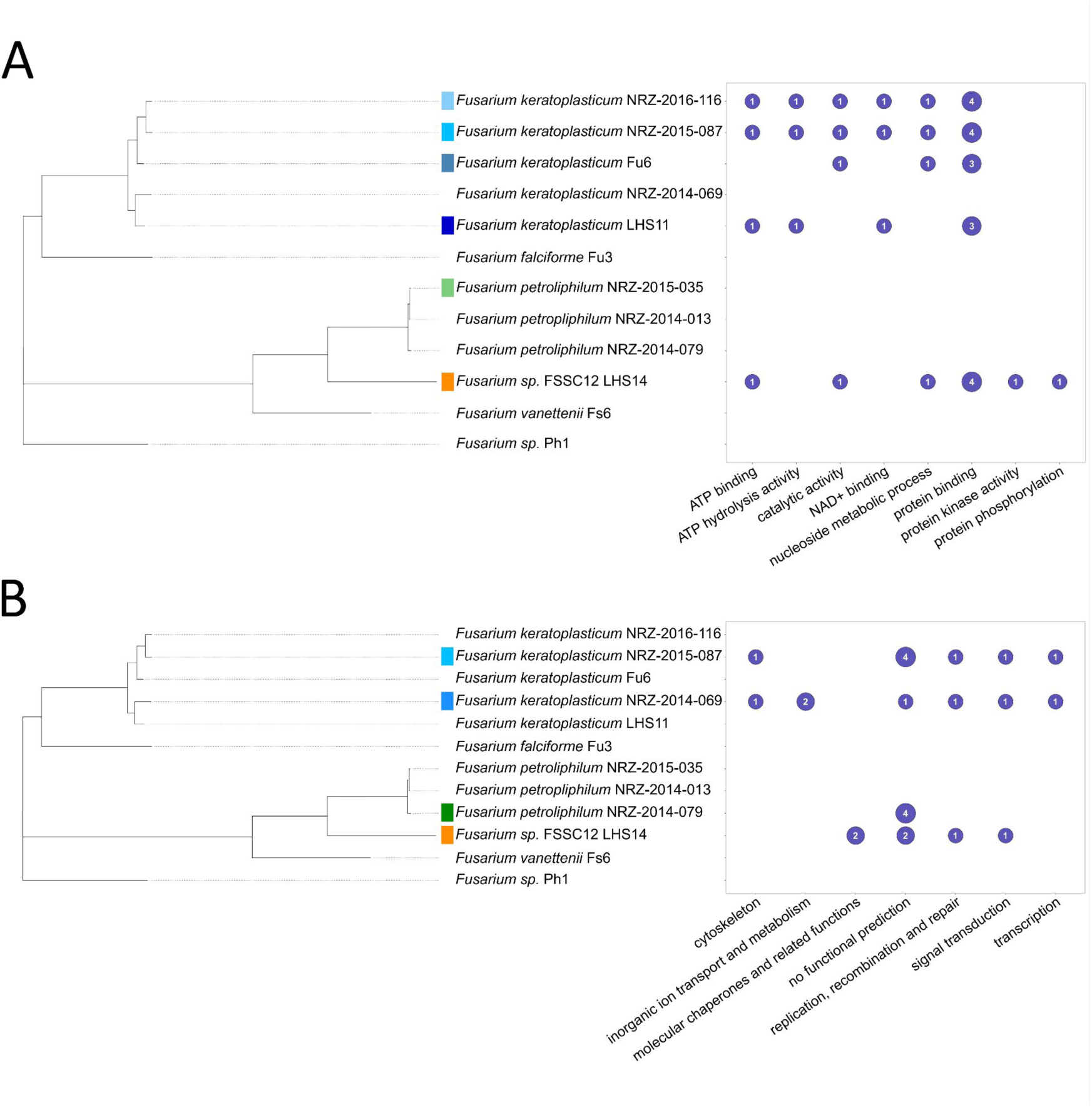
Functional annotations of cargo of *Arwing* and *Galactica Starships*. (a) GO biological process terms and their gene copy number in *Arwing* cargo. (b) COG annotations and their copy number in *Galactica Starships*.

**Supplementary Figure 8.**
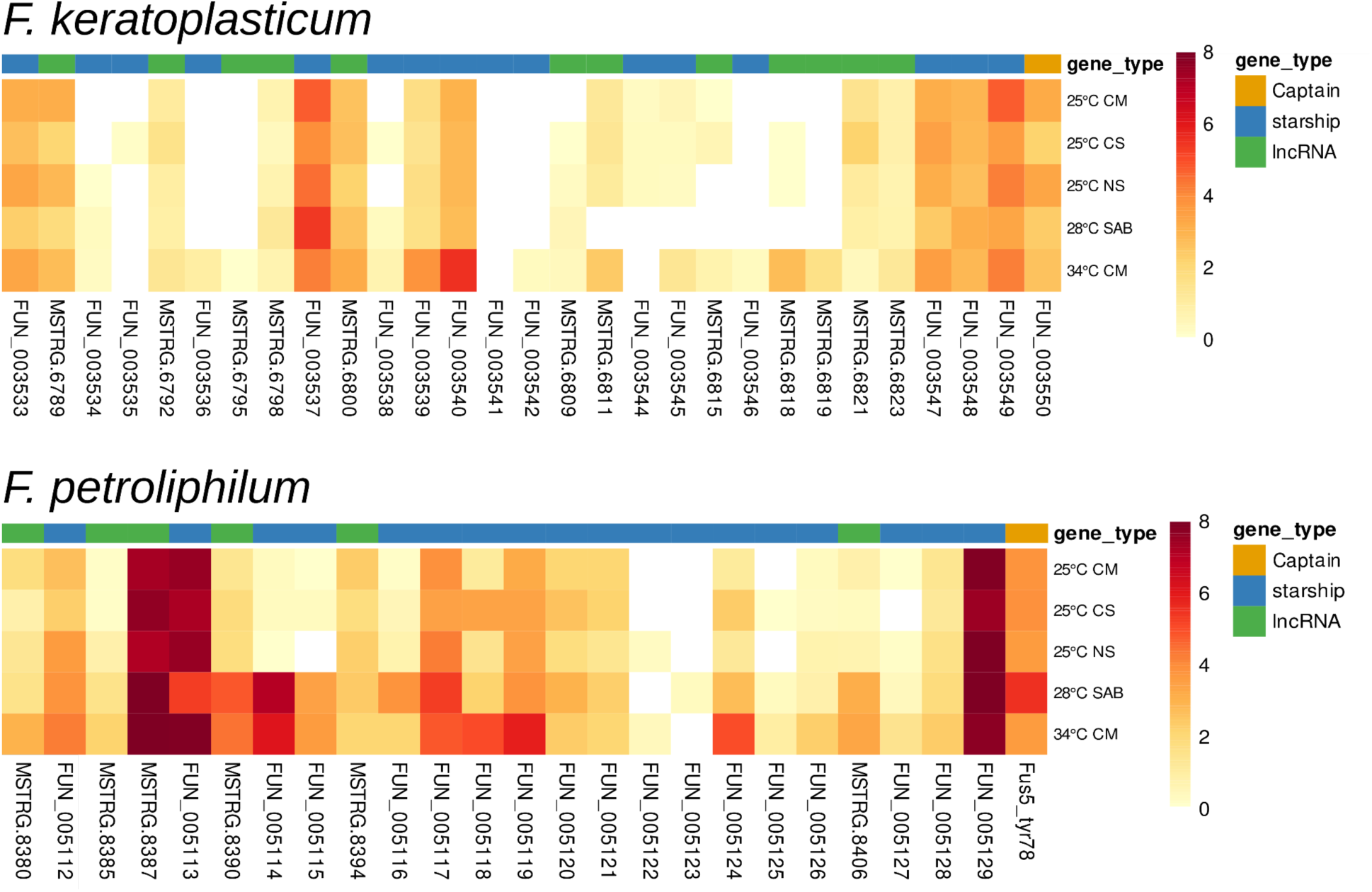
Expression of *Galactica Starship* cargo genes and associated lncRNAs in *F. keratoplasticum* NRZ-2014-069 and *F. petroliphilum* NRZ-2014-079 across different culture conditions. The heat map displays log₂-transformed TPM values (log₂(TPM+1)) for genes within the *Starship* cluster and co-localized lncRNAs under five experimental conditions: 25°C complete medium (CM), 25°C carbon starvation (CS), 25°C nitrogen starvation (NS), 34°C complete medium (CM), and 28°C conidia grown on solid media (SAB). Each value represents the average expression of two biological replicates. Genes are ordered according to their genomic position in the *Starship* element. Gene types are color-coded in the annotation bar: orange indicates the captain gene, blue represents *Starship* genes, and green denotes lincRNAs. Genes ordered by synteny.

**Supplementary Figure 9.**
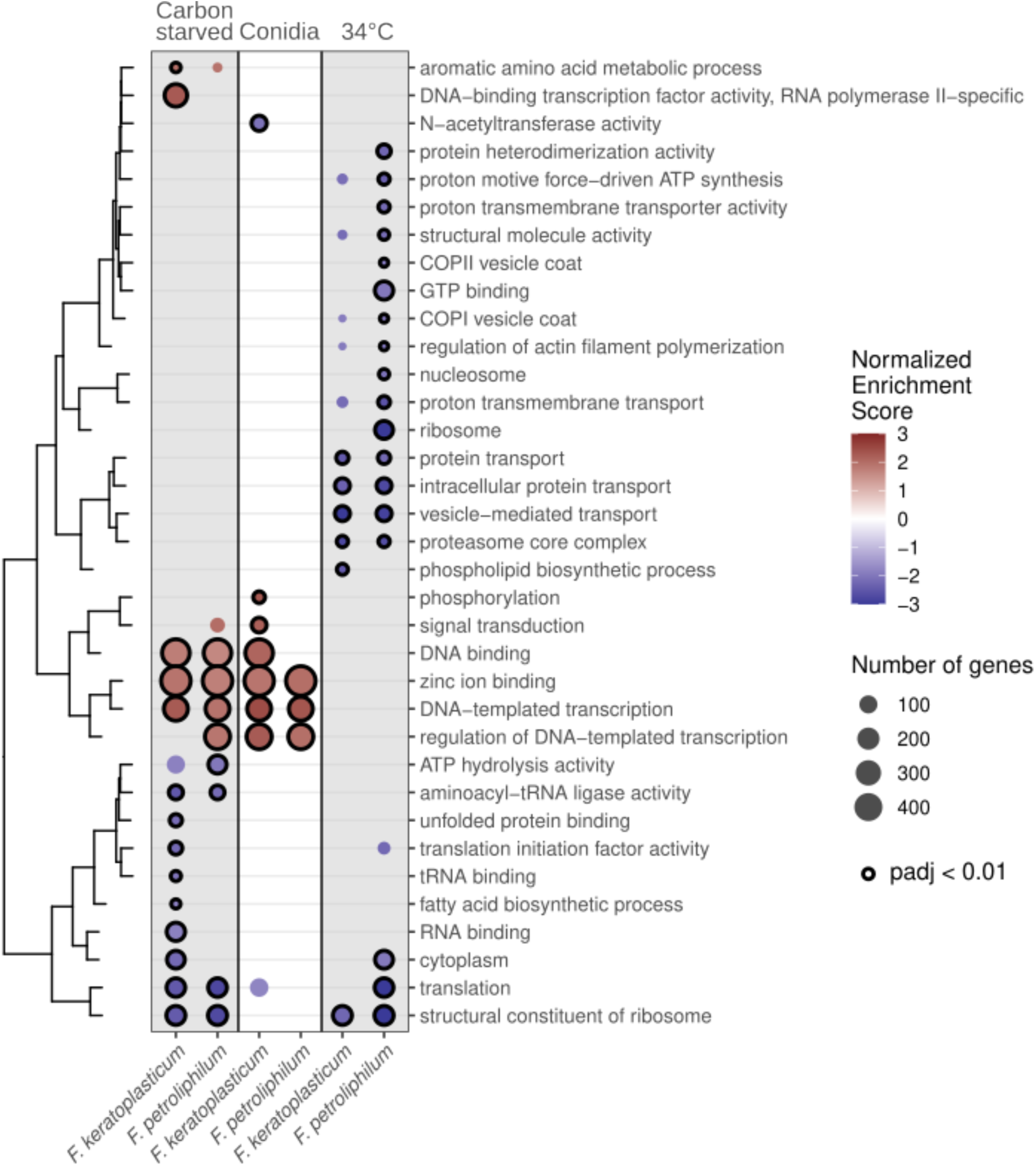
Comparison of enriched and depleted Gene Ontology terms in the transcriptomes of *F. keratoplasticum* and *F. petroliphilum*. Experiments were conducted using NRZ-2014-069 for *F. keratoplasticum* and NRZ-2014-079 for *F. petroliphilum*. Differential expression for the conditions shown was calculated in comparison to mycelia grown in complete media at 25°C. Color indicates normalized enrichment score calculated by fgsea. Circle size indicates the number of genes for each GO term that were differentially expressed. Circles with a black outline indicate GO terms for which adjusted p-value < 0.01. All other circles have an adjusted p-value < 0.1.

## Supplementary File List

Supplementary File 1. Accession numbers for the genome assemblies used in the phylogeny, including the publicly available FSSC isolates analyzed.

Supplementary File 2. Gene Ontology terms significantly over- or underrepresented in the genomes of *F. keratoplasticum* and *F. petroliphilum*. File includes the gene counts and z-scored values depicted in Figure 5. Description of genome-wide ontology terms.

Supplementary File 3. Conservation of lncRNAs between *F. keratoplasticum* and *F. petroliphilum* identified through nucleotide alignment of lncRNA sequences and synteny using flanking protein-coding genes.

Supplementary File 4. Differentially expressed genes in conidia, under carbon starvation, and at 34°C relative to mycelia grown in complete media. Description of DEG file.

Supplementary File 5. Gene set enrichment analysis of differentially expressed genes in *F. keratoplasticum* and *F. petroliphilum*.

Supplementary File 6. Newick phylogeny displayed in Figure 1.

Supplementary File 7. Core and accessory genes for the FSSC, *F. keratoplasticum* and *F. petroliphilum*

Supplementary File 8. Characterization of lncRNA-BGC regulatory networks and expression profiles. Summary of lncRNA-BGC co-expression interactions in *F. keratoplasticum* and *F. petroliphilum*, including the number of connected clusters and the identity of biosynthetic products. Transcriptional activity metrics for lncRNAs and BGC backbone enzymes across five growth conditions.

## References

1. Coleman JJ. 2016. The Fusarium solani species complex: Ubiquitous pathogens of agricultural importance. Molecular Plant Pathology 17:146–158.

2. Singh RP, Singh PK, Rutkoski J, Hodson DP, He X, Jørgensen LN, Hovmøller MS, Huerta-Espino J. 2016. Disease Impact on Wheat Yield Potential and Prospects of Genetic Control. Annu Rev Phytopathol 54:303–322.

3. Goswami RS, Kistler HC. 2004. Heading for disaster: Fusarium graminearum on cereal crops. Molecular Plant Pathology 5:515–525.

4. Gordon TR. 2017. Fusarium oxysporum and the Fusarium Wilt Syndrome. Annual Review of Phytopathology 55:23–39.

5. Dean R, Van Kan JAL, Pretorius ZA, Hammond-Kosack KE, Di Pietro A, Spanu PD, Rudd JJ, Dickman M, Kahmann R, Ellis J, Foster GD. 2012. The Top 10 fungal pathogens in molecular plant pathology. Mol Plant Pathol 13:414–430.

6. Nelson PE, Dignani MC, Anaissie EJ. 1994. Taxonomy, biology, and clinical aspects of Fusarium species. Clinical Microbiology Reviews 7:479–504.

7. Nucci M, Anaissie E. 2023. Invasive fusariosis. Clinical Microbiology Reviews 36:e00159–22.

8. Walther G, Stasch S, Kaerger K, Hamprecht A, Roth M, Cornely OA, Geerling G, Mackenzie CR, Kurzai O, von Lilienfeld-Toal M. 2017. Fusarium Keratitis in Germany. J Clin Microbiol.

9. Cabañes FJ, Alonso JM, Castellá G, Alegre F, Domingo M, Pont S. 1997. Cutaneous hyalohyphomycosis caused by Fusarium solani in a loggerhead sea turtle (Caretta caretta L.). J Clin Microbiol 35:3343–3345.

10. Gleason FH, Allerstorfer M, Lilje O. Newly emerging diseases of marine turtles, especially sea turtle egg fusariosis (SEFT), caused by species in the Fusarium solani complex (FSSC). Mycology 11:184–194.

11. Hoh DZ, Lee H-H, Wada N, Liu W-A, Lu MR, Lai C-K, Ke H-M, Sun P-F, Tang S-L, Chung W-H, Chen Y-L, Chung C-L, Tsai IJ. 2022. Comparative genomic and transcriptomic analyses of trans-kingdom pathogen Fusarium solani species complex reveal degrees of compartmentalization. BMC Biol 20:1–18.

12. Batista BG, de Chaves MA, Reginatto P, Saraiva OJ, Fuentefria AM. 2020. Human fusariosis: An emerging infection that is difficult to treat. Rev Soc Bras Med Trop 53.

13. World Heath Organization. 2022. WHO fungal priority pathogens list to guide research, development and public health action. World Health Organization.

14. Campo M, Lewis RE, Kontoyiannis DP. 2010. Invasive fusariosis in patients with hematologic malignancies at a cancer center: 1998-2009. J Infect 60:331–337.

15. Demonchy J, Biard L, Clere-Jehl R, Wallet F, Mokart D, Moreau A-S, Argaud L, Verlhac C, Pène F, Lautrette A, Bige N, de Jong A, Canet E, Quenot J-P, Issa N, Zerbib Y, Bouard I, Picard M, Zafrani L. 2024. Multicenter Retrospective Study of Invasive Fusariosis in Intensive Care Units, France. Emerging Infectious Diseases 10.3201/eid3002.231221.

16. Nucci M, Marr KA, Vehreschild MJGT, de Souza CA, Velasco E, Cappellano P, Carlesse F, Queiroz-Telles F, Sheppard DC, Kindo A, Cesaro S, Hamerschlak N, Solza C, Heinz WJ, Schaller M, Atalla A, Arikan-Akdagli S, Bertz H, Galvão Castro C, Herbrecht R, Hoenigl M, Härter G, Hermansen NEU, Josting A, Pagano L, Salles MJC, Mossad SB, Ogunc D, Pasqualotto AC, Araujo V, Troke PF, Lortholary O, Cornely OA, Anaissie E. 2014. Improvement in the outcome of invasive fusariosis in the last decade. Clin Microbiol Infect 20:580–585.

17. Horn DL, Freifeld AG, Schuster MG, Azie NE, Franks B, Kauffman CA. 2014. Treatment and outcomes of invasive fusariosis: review of 65 cases from the PATH Alliance(®) registry. Mycoses 57:652–658.

18. van der Does Ma L-J HC, Borkovich KA, Coleman JJ, Daboussi M-J, Di Pietro A, Dufresne M, Freitag M, Grabherr M, Henrissat B, Houterman PM, Kang S, Shim W-B, Woloshuk C, Xie X, Xu J-R, Antoniw J, Baker SE, Bluhm BH, Breakspear A, Brown DW, Butchko RAE, Chapman S, Coulson R, Coutinho PM, Danchin EGJ, Diener A, Gale LR, Gardiner DM, Goff S, Hammond-Kosack KE, Hilburn K, Hua-Van A, Jonkers W, Kazan K, Kodira CD, Koehrsen M, Kumar L, Lee Y-H, Li L, Manners JM, Miranda-Saavedra D, Mukherjee M, Park G, Park J, Park S-Y, Proctor RH, Regev A, Ruiz-Roldan MC, Sain D, Sakthikumar S, Sykes S, Schwartz DC, Turgeon BG, Wapinski I, Yoder O, Young S, Zeng Q, Zhou S, Galagan J, Cuomo CA, Kistler HC, Rep M. 2010. Comparative genomics reveals mobile pathogenicity chromosomes in Fusarium. Nature 464:367–373.

19. Zhang Y, Yang H, Turra D, Zhou S, Ayhan DH, DeIulio GA, Guo L, Broz K, Wiederhold N, Coleman JJ, Donnell KO, Youngster I, McAdam AJ, Savinov S, Shea T, Young S, Zeng Q, Rep M, Pearlman E, Schwartz DC, Di Pietro A, Kistler HC, Ma LJ. 2020. The genome of opportunistic fungal pathogen Fusarium oxysporum carries a unique set of lineage-specific chromosomes. Communications Biology 3:1–12.

20. 2019. Genome-wide analysis of Fusarium verticillioides reveals inter-kingdom contribution of horizontal gene transfer to the expansion of metabolism. Fungal Genet Biol 128:60–73.

21. Guarro J. 2013. Fusariosis, a complex infection caused by a high diversity of fungal species refractory to treatment. Eur J Clin Microbiol Infect Dis 32:1491–1500.

22. van Diepeningen AD, Al-Hatmi AMS, Brankovics B, de Hoog GS. 2014. Taxonomy and Clinical Spectra of Fusarium Species: Where Do We Stand in 2014? Current Clinical Microbiology Reports 1:10–18.

23. Short DPG, O’Donnell K, Zhang N, Juba JH, Geiser DM. 2011. Widespread occurrence of diverse human pathogenic types of the fungus Fusarium detected in plumbing drains. J Clin Microbiol 49:4264–4272.

24. Garnica M, Nucci M. 2013. Epidemiology of Fusariosis. Curr Fungal Infect Rep 7:301–305.

25. Zhang N, O’Donnell K, Sutton DA, Nalim FA, Summerbell RC, Padhye AA, Geiser DM. 2006. Members of the Fusarium solani Species Complex That Cause Infections in Both Humans and Plants Are Common in the Environment. Journal of Clinical Microbiology 44:2186–2190.

26. Hoenigl M, Jenks JD, Egger M, Nucci M, Thompson GR. 2023. Treatment of Fusarium Infection of the Central Nervous System: A Review of Past Cases to Guide Therapy for the Ongoing 2023 Outbreak in the United States and Mexico. Mycopathologia 188:973–981.

27. Strong N, Meeks G, Sheth SA, McCullough L, Villalba JA, Tan C, Barreto A, Wanger A, McDonald M, Kan P, Shaltoni H, Maldonado JC, Parada V, Hassan AE, Reagan-Steiner S, Chiller T, Gold JAW, Smith DJ, Ostrosky-Zeichner L. 2024. Neurovascular Complications of Iatrogenic Fusarium solani Meningitis. New England Journal of Medicine 390:522–529.

28. García-Rodríguez G, Duque-Molina C, Kondo-Padilla I, Zaragoza-Jiménez CA, González-Cortés VB, Flores-Antonio R, Villa-Reyes T, Vargas-Rubalcava A, Ruano-Calderon LÁ, Tinoco-Favila JC, Sánchez-Salazar HC, Rivas-Ruiz R, Castro-Escamilla O, Martínez-Gamboa RA, González-Lara F, López-Martínez I, Chiller TM, Pelayo R, Bonifaz LC, Robledo-Aburto Z, Alcocer-Varela J. 2024. Outbreak of Meningitis in Immunocompetent Persons Associated With Neuraxial Blockade in Durango, Mexico, 2022-2023. Open Forum Infect Dis 11:ofad690.

29. Gibert S, Edel-Hermann V, Gautheron E, Gautheron N, Sol J-M, Capelle G, Galland R, Bardon-Debats A, Lambert C, Steinberg C. 2022. First Report of Fusarium avenaceum, Fusarium oxysporum, Fusarium redolens, and Fusarium solani Causing Root Rot in Pea in France. Plant Dis.

30. Pérez-Hernández A, Porcel-Rodríguez E, Gómez-Vázquez J. 2017. Survival of Fusarium solani f. sp. cucurbitae and Fungicide Application, Soil Solarization, and Biosolarization for Control of Crown and Foot Rot of Zucchini Squash. Plant Dis.

31. Gómez J, Serrano Y, Pérez A, Porcel E, Gómez R, Aguilar MI. 2014. Fusarium solani f. sp. cucurbitae, affecting melon in Almería Province, Spain. Australas Plant Dis Notes 9:1–3.

32. Wang R-Y, Gao B, Li X-H, Ma J, Chen S-L. 2013. First Report of Fusarium solani Causing Fusarium Root Rot and Stem Canker on Storage Roots of Sweet Potato in China.

33. Vega-Gutiérrez TA, López-Orona CA, Molina-Cárdenas L, López-Urquídez GA, Payán-Arzapalo MA, Tirado-Ramírez MA. 2023. First Report of Fusarium keratoplasticum Causing Strawberry Root Rot in Sinaloa, Mexico. Plant Dis.

34. Rosa, P. D., Heidrich, D., Corrêa, C., Scroferneker, M. L., Vettorato, G., Fuentefria, A. M., & Goldani, L. Z. 2017. Genetic diversity and antifungal susceptibility of Fusarium isolates in onychomycosis. Mycoses 60:616–622.

35. Peinado-Acevedo JS, Ramírez-Sánchez IC. 2020. Endocarditis by Fusarium keratoplasticum. Mycopathologia 186:131–133.

36. Short DPG, O’Donnell K, Thrane U, Nielsen KF, Zhang N, Juba JH, Geiser DM. 2013. Phylogenetic relationships among members of the Fusarium solani species complex in human infections and the descriptions of F. keratoplasticum sp. nov. and F. petroliphilum stat. nov. Fungal Genetics and Biology 53:59–70.

37. Mattick JS, Amaral PP, Carninci P, Carpenter S, Chang HY, Chen L-L, Chen R, Dean C, Dinger ME, Fitzgerald KA, Gingeras TR, Guttman M, Hirose T, Huarte M, Johnson R, Kanduri C, Kapranov P, Lawrence JB, Lee JT, Mendell JT, Mercer TR, Moore KJ, Nakagawa S, Rinn JL, Spector DL, Ulitsky I, Wan Y, Wilusz JE, Wu M. 2023. Long non-coding RNAs: definitions, functions, challenges and recommendations. Nat Rev Mol Cell Biol 24:430–447.

38. Kim W, Miguel-Rojas C, Wang J, Townsend JP, Trail F. 2018. Developmental Dynamics of Long Noncoding RNA Expression during Sexual Fruiting Body Formation in Fusarium graminearum. MBio 9.

39. Parra-Rivero O, Pardo-Medina J, Gutiérrez G, Limón MC, Avalos J. 2020. A novel lncRNA as a positive regulator of carotenoid biosynthesis in Fusarium. Sci Rep 10:1–14.

40. Wang J, Zeng W, Cheng J, Xie J, Fu Y, Jiang D, Lin Y. 2021. lncRsp1, a long noncoding RNA, influences Fgsp1 expression and sexual reproduction in Fusarium graminearum. Mol Plant Pathol 23:265–277.

41. Kingsbury JM, Yang Z, Ganous TM, Cox GM, McCusker JH. 2004. Cryptococcus neoformans Ilv2p confers resistance to sulfometuron methyl and is required for survival at 37 °C and in vivo. Microbiology 150:1547–1558.

42. Garcia MD, Chua SMH, Low Y-S, Lee Y-T, Agnew-Francis K, Wang J-G, Nouwens A, Lonhienne T, Williams CM, Fraser JA, Guddat LW. 2018. Commercial AHAS-inhibiting herbicides are promising drug leads for the treatment of human fungal pathogenic infections. Proceedings of the National Academy of Sciences 115:E9649–E9658.

43. Oh Y, Franck WL, Han S-O, Shows A, Gokce E, Muddiman DC, Dean RA. 2012. Polyubiquitin Is Required for Growth, Development and Pathogenicity in the Rice Blast Fungus Magnaporthe oryzae. PLoS One 7.

44. Rodier M-H, El Moudni B, Kauffmann-Lacroix C, Daniault G, Jacquemin J-L. 1999. A Candida albicans metallopeptidase degrades constitutive proteins of extracellular matrix. FEMS Microbiol Lett 177:205–210.

45. Silva BA, Souza-Gonçalves AL, Pinto MR, Barreto-Bergter E, Santos ALS. 2011. Metallopeptidase inhibitors arrest vital biological processes in the fungal pathogen Scedosporium apiospermum. Mycoses 54:105–112.

46. King BC, Waxman KD, Nenni NV, Walker LP, Bergstrom GC, Gibson DM. 2011. Arsenal of plant cell wall degrading enzymes reflects host preference among plant pathogenic fungi. Biotechnol Biofuels 4:4.

47. Skamnioti P, Gurr SJ. 2007. Magnaporthe grisea Cutinase2 Mediates Appressorium Differentiation and Host Penetration and Is Required for Full Virulence. Plant Cell 19:2674–2689.

48. Rogers LM, Flaishman MA, Kolattukudy PE. 1994. Cutinase gene disruption in Fusarium solani f sp pisi decreases its virulence on pea. Plant Cell 6:935–945.

49. O’Brien J, Wright GD. 2011. An ecological perspective of microbial secondary metabolism. Current Opinion in Biotechnology 22:552–558.

50. Swayambhu G, Bruno M, Gulick AM, Pfeifer BA. 2021. Siderophore natural products as pharmaceutical agents. Curr Opin Biotechnol 69:242–251.

51. Dufour N, Rao RP. 2011. Secondary metabolites and other small molecules as intercellular pathogenic signals. FEMS Microbiol Lett 314:10–17.

52. Jihua Wei BW. 2020. Chemistry and bioactivities of secondary metabolites from the genus Fusarium. Fitoterapia 146:104638.

53. Li M, Yu R, Bai X, Wang H, Zhang H. 2020. Fusarium: a treasure trove of bioactive secondary metabolites. Nat Prod Rep 37:1568–1588.

54. Dawson MJ, Farthing JE, Marshall PS, Middleton RF, O’neill MJ, Shuttleworth A, Stylli C, Murray Tait R, Taylor PM, Wildman HG, Buss AD, Langley D, Hayes MV. 1992. THE SQUALESTATINS, NOVEL INHIBITORS OF SQUALENE SYNTHASE PRODUCED BY A SPECIES OF PHOMA I. TAXONOMY, FERMENTATION, ISOLATION, PHYSICO-CHEMICAL PROPERTIES AND BIOLOGICAL ACTIVITY. J Antibiot 45:639–647.

55. Kurobane I, Vining LC, Gavin Mcinnes A, Gerber NN. 1980. METABOLITES OF FUSARIUM SOLANI RELATED TO DIHYDROFUSARUBIN. J Antibiot 33:1376–1379.

56. Kundu A, Mandal A, Saha S, Prabhakaran P, Walia S. 2020. Fungicidal activity and molecular modeling of fusarubin analogues from Fusarium oxysporum. Toxicol Environ Chem.

57. Costa JH, Wassano CI, Angolini CFF, Scherlach K, Hertweck C, Pacheco Fill T. 2019. Antifungal potential of secondary metabolites involved in the interaction between citrus pathogens. Sci Rep 9:1–11.

58. Kato S, Motoyama T, Futamura Y, Uramoto M, Nogawa T, Hayashi T, Hirota H, Tanaka A, Takahashi-Ando N, Kamakura T, Osada H. 2020. Biosynthetic gene cluster identification and biological activity of lucilactaene from Fusarium sp. RK97-94. Biosci Biotechnol Biochem 84:1303–1307.

59. Suga H, Arai M, Fukasawa E, Motohashi K, Nakagawa H, Tateishi H, Fuji S-I, Shimizu M, Kageyama K, Hyakumachi M. 2018. Genetic Differentiation Associated with Fumonisin and Gibberellin Production in Japanese Fusarium fujikuroi. Appl Environ Microbiol.

60. Hossain MT, Khan A, Chung EJ, Rashid MH-O, Chung YR. 2016. Biological Control of Rice Bakanae by an Endophytic Bacillus oryzicola YC7007. Plant Pathol J 32:228–241.

61. Urquhart AS, Gluck-Thaler E, Vogan AA. 2024. Gene acquisition by giant transposons primes eukaryotes for rapid evolution via horizontal gene transfer. Science Advances 10:eadp8738.

62. Urquhart AS, O’Donnell S, Gluck-Thaler E, Vogan AA. 2025. A natural mechanism of eukaryotic horizontal gene transfer. bioRxiv 2025.02.28.640899.

63. Bucknell A, Wilson HM, Gonçalves Dos Santos KC, Simpfendorfer S, Milgate A, Germain H, Solomon PS, Bentham A, McDonald MC. 2025. Sanctuary: a Starship transposon facilitating the movement of the virulence factor ToxA in fungal wheat pathogens. mBio 16:e0137125.

64. Urquhart A, Vogan AA, Gluck-Thaler E. 2024. Starships: a new frontier for fungal biology. Trends in Genetics 40:1060–1073.

65. Sato Y, Bex R, van den Berg GCM, Santhanam P, Höfte M, Seidl MF, Thomma BPHJ. 2025. Starship giant transposons dominate plastic genomic regions in a fungal plant pathogen and drive virulence evolution. Nat Commun 16:6806.

66. Peck LD, Llewellyn T, Bennetot B, O’Donnell S, Nowell RW, Ryan MJ, Flood J, de la Vega RCR, Ropars J, Giraud T, Spanu PD, Barraclough TG. 2024. Horizontal transfers between fungal Fusarium species contributed to successive outbreaks of coffee wilt disease. PLOS Biology 22:e3002480.

67. Hughes TR, Marton MJ, Jones AR, Roberts CJ, Stoughton R, Armour CD, Bennett HA, Coffey E, Dai H, Et R-M, He YD, Kidd MJ, King AM, Meyer MR, Slade D, Lum PY, Stepaniants SB, Shoemaker DD. 2000. Functional Discovery via a Compendium of Expression Profiles they screen for phenotypes. However, just as high. Cell 102:109–126.

68. Eisen MB, Spellman PT, Brown PO, Botstein D. 1998. Cluster analysis and display of genome-wide expression patterns. Proceedings of the National Academy of Sciences 95:14863–14868.

69. Fayyaz A, Robinson G, Chang PL, Bekele D, Yimer S, Carrasquilla-Garcia N, Negash K, Surendrarao A, von Wettberg EJB, Kemal S-A, Tesfaye K, Fikre A, Farmer AD, Cook DR. 2023. Hiding in plain sight: Genome-wide recombination and a dynamic accessory genome drive diversity in Fusarium oxysporum f.sp. ciceris. Proceedings of the National Academy of Sciences 120:e2220570120.

70. van Westerhoven AC, Aguilera-Galvez C, Nakasato-Tagami G, Shi-Kunne X, Martinez de la Parte E, Chavarro-Carrero E, Meijer HJG, Feurtey A, Maryani N, Ordóñez N, Schneiders H, Nijbroek K, Wittenberg AHJ, Hofstede R, García-Bastidas F, Sørensen A, Swennen R, Drenth A, Stukenbrock EH, Kema GHJ, Seidl MF. Segmental duplications drive the evolution of accessory regions in a major crop pathogen. New Phytologist n/a.

71. Barber AE, Sae-ong T, Kang K, Seelbinder B, Li J, Walther G, Panagiotou G, Kurzai O. 2021. Aspergillus fumigatus pan-genome analysis identifies genetic variants associated with human infection. Nature Microbiology 6.

72. McCarthy CGP, Fitzpatrick DA. 2019. Pan-genome analyses of model fungal species. Microbial Genomics 1–23.

73. Gangurde SS, Korani W, Bajaj P, Wang H, Fountain JC, Agarwal G, Pandey MK, Abbas HK, Chang P-K, Holbrook CC, Kemerait RC, Varshney RK, Dutta B, Clevenger JP, Guo B. 2024. Aspergillus flavus pangenome (AflaPan) uncovers novel aflatoxin and secondary metabolite associated gene clusters. BMC Plant Biology 24:354.

74. Hatmaker EA, Barber AE, Drott MT, Sauters TJC, Gumilang A, Alastruey-Izquierdo A, Garcia-Hermoso D, Eagan JL, Keller NP, Kontoyiannis DP, Kurzai O, Rokas A. 2025. Population structure in a fungal human pathogen is potentially linked to pathogenicity. Nat Commun 16:7594.

75. Dewar AE, Hao C, Belcher LJ, Ghoul M, West SA. 2024. Bacterial lifestyle shapes pangenomes. Proceedings of the National Academy of Sciences 121:e2320170121.

76. Perrier M, Barber AE. 2024. Unraveling the genomic diversity and virulence of human fungal pathogens through pangenomics. PLoS Pathog 20:e1012313.

77. Brůna T, Sreedasyam A, Harder AM, Lovell JT. 2026. Evolutionary and methodological considerations when interpreting gene presence–absence variation in pangenomes. NAR Genom Bioinform 8:lqag011.

78. Möller M, Stukenbrock EH. 2017. Evolution and genome architecture in fungal plant pathogens. Nat Rev Microbiol 15:756–771.

79. Fokkens L, Guo L, Dora S, Wang B, Ye K, Sánchez-Rodríguez C, Croll D. 2020. A Chromosome-Scale Genome Assembly for the Fusarium oxysporum Strain Fo5176 To Establish a Model Arabidopsis-Fungal Pathosystem. G3 Genes|Genomes|Genetics 10:3549–3555.

80. Hovhannisyan H, Gabaldón T. 2021. The long non-coding RNA landscape of Candida yeast pathogens. Nat Commun 12.

81. Kalem MC, Panepinto JC. 2022. Long Non-Coding RNAs in Cryptococcus neoformans: Insights Into Fungal Pathogenesis. Front Cell Infect Microbiol 12:858317.

82. Glad HM, Tralamazza SM, Croll D. 2023. The expression landscape and pangenome of long non-coding RNA in the fungal wheat pathogen Zymoseptoria tritici. Microbial Genomics 9:001136.

83. Gao J, Chow EWL, Wang H, Xu X, Cai C, Song Y, Wang J, Wang Y. 2021. LncRNA DINOR is a virulence factor and global regulator of stress responses in Candida auris. Nat Microbiol 6:842–851.

84. Kolmogorov M, Yuan J, Lin Y, Pevzner PA. 2019. Assembly of long, error-prone reads using repeat graphs. 5. Nat Biotechnol 37:540–546.

85. Walker BJ, Abeel T, Shea T, Priest M, Abouelliel A, Sakthikumar S, Cuomo CA, Zeng Q, Wortman J, Young SK, Earl AM. 2014. Pilon: An Integrated Tool for Comprehensive Microbial Variant Detection and Genome Assembly Improvement. PLOS ONE 9:e112963.

86. Alonge M, Lebeigle L, Kirsche M, Jenike K, Ou S, Aganezov S, Wang X, Lippman ZB, Schatz MC, Soyk S. 2022. Automated assembly scaffolding using RagTag elevates a new tomato system for high-throughput genome editing. Genome Biol 23:258.

87. Astashyn A, Tvedte ES, Sweeney D, Sapojnikov V, Bouk N, Joukov V, Mozes E, Strope PK, Sylla PM, Wagner L, Bidwell SL, Brown LC, Clark K, Davis EW, Smith-White B, Hlavina W, Pruitt KD, Schneider VA, Murphy TD. 2024. Rapid and sensitive detection of genome contamination at scale with FCS-GX. Genome Biol 25:60.

88. Flynn JM, Hubley R, Goubert C, Rosen J, Clark AG, Feschotte C, Smit AF. 2020. RepeatModeler2 for automated genomic discovery of transposable element families. Proceedings of the National Academy of Sciences 117:9451–9457.

89. Tarailo-Graovac M, Chen N. 2009. Using RepeatMasker to identify repetitive elements in genomic sequences. Curr Protoc Bioinformatics Chapter 4:4.10.1-4.10.14.

90. Pertea M, Pertea GM, Antonescu CM, Chang T-C, Mendell JT, Salzberg SL. 2015. StringTie enables improved reconstruction of a transcriptome from RNA-seq reads. Nat Biotechnol 33:290–295.

91. Sims D, Ilott NE, Sansom SN, Sudbery IM, Johnson JS, Fawcett KA, Berlanga-Taylor AJ, Luna-Valero S, Ponting CP, Heger A. 2014. CGAT: computational genomics analysis toolkit. Bioinformatics 30:1290–1291.

92. Kang Y-J, Yang D-C, Kong L, Hou M, Meng Y-Q, Wei L, Gao G. 2017. CPC2: a fast and accurate coding potential calculator based on sequence intrinsic features. Nucleic Acids Res 45:W12–W16.

93. Wucher V, Legeai F, Hédan B, Rizk G, Lagoutte L, Leeb T, Jagannathan V, Cadieu E, David A, Lohi H, Cirera S, Fredholm M, Botherel N, Leegwater PAJ, Le Béguec C, Fieten H, Johnson J, Alföldi J, André C, Lindblad-Toh K, Hitte C, Derrien T. 2017. FEELnc: a tool for long non-coding RNA annotation and its application to the dog transcriptome. Nucleic Acids Res 45:e57.

94. Liao Y, Smyth GK, Shi W. 2014. featureCounts: an efficient general purpose program for assigning sequence reads to genomic features. Bioinformatics 30:923–930.

95. Emms DM, Kelly S. 2019. OrthoFinder: phylogenetic orthology inference for comparative genomics. Genome Biol 20:238.

96. Edgar RC. 2004. MUSCLE: multiple sequence alignment with high accuracy and high throughput. Nucleic Acids Res 32:1792–1797.

97. Steenwyk JL, Iii TJB, Li Y, Shen X-X, Rokas A. 2020. ClipKIT: A multiple sequence alignment trimming software for accurate phylogenomic inference. PLOS Biology 18:e3001007.

98. Minh BQ, Schmidt HA, Chernomor O, Schrempf D, Woodhams MD, von Haeseler A, Lanfear R. 2020. IQ-TREE 2: New Models and Efficient Methods for Phylogenetic Inference in the Genomic Era. Molecular Biology and Evolution 37:1530–1534.

99. Lovell JT, Sreedasyam A, Schranz ME, Wilson M, Carlson JW, Harkess A, Emms D, Goodstein DM, Schmutz J. 2022. GENESPACE tracks regions of interest and gene copy number variation across multiple genomes. eLife 11:e78526.

100. Marçais G, Delcher AL, Phillippy AM, Coston R, Salzberg SL, Zimin A. 2018. MUMmer4: A fast and versatile genome alignment system. PLOS Computational Biology 14:e1005944.

101. Pegueroles C, Iraola-Guzmán S, Chorostecki U, Ksiezopolska E, Saus E, Gabaldón T. 2019. Transcriptomic analyses reveal groups of co-expressed, syntenic lncRNAs in four species of the genus Caenorhabditis. RNA Biol 16:320–329.

102. Chen S, Zhou Y, Chen Y, Gu J. 2018. fastp: an ultra-fast all-in-one FASTQ preprocessor. Bioinformatics 34:i884–i890.

103. Dobin A, Davis CA, Schlesinger F, Drenkow J, Zaleski C, Jha S, Batut P, Chaisson M, Gingeras TR. 2013. STAR: ultrafast universal RNA-seq aligner. Bioinformatics 29:15–21.

104. Love MI, Huber W, Anders S. 2014. Moderated estimation of fold change and dispersion for RNA-seq data with DESeq2. Genome Biol 15:550.

105. Korotkevich G, Sukhov V, Budin N, Shpak B, Artyomov MN, Sergushichev A. 2021. Fast gene set enrichment analysis. bioRxiv 060012.

106. Blin K, Shaw S, Augustijn HE, Reitz ZL, Biermann F, Alanjary M, Fetter A, Terlouw BR, Metcalf WW, Helfrich EJN, van Wezel GP, Medema MH, Weber T. 2023. antiSMASH 7.0: new and improved predictions for detection, regulation, chemical structures and visualisation. Nucleic Acids Res 51:W46–W50.

107. Navarro-Muñoz JC, Selem-Mojica N, Mullowney MW, Kautsar SA, Tryon JH, Parkinson EI, De Los Santos ELC, Yeong M, Cruz-Morales P, Abubucker S, Roeters A, Lokhorst W, Fernandez-Guerra A, Cappelini LTD, Goering AW, Thomson RJ, Metcalf WW, Kelleher NL, Barona-Gomez F, Medema MH. 2020. A computational framework to explore large-scale biosynthetic diversity. Nat Chem Biol 16:60–68.

108. Gluck-Thaler E, Vogan AA. 2024. Systematic identification of cargo-mobilizing genetic elements reveals new dimensions of eukaryotic diversity. Nucleic Acids Res 52:5496–5513.

109. Li H. 2018. Minimap2: pairwise alignment for nucleotide sequences. Bioinformatics 34:3094–3100.

110. Langfelder P, Horvath S. 2008. WGCNA: an R package for weighted correlation network analysis. BMC Bioinformatics 9:559.

